# Systemic multi-omic remodelling underlies health benefits of intermittent fasting

**DOI:** 10.1101/2025.08.30.673138

**Authors:** Chiara Maria Stella Herzog, Charlotte D. Vavourakis, Bente Theeuwes, Elisa Redl, Christina Watschinger, Gabriel Knoll, Magdalena Hagen, Andreas Haider, Hans-Peter Platzer, Umesh Kumar, Sophia Zollner-Kiechl, Maria Cavinato, Pidder Jansen-Duerr, Matthias Schmuth, Maximilian Lammer, Michael Knoflach, Verena Lindner, Anna Wimmer, Peter Widschwendter, Tobias Greitemeyer, Sonja Sturm, Hermann Stuppner, Birgit Weinberger, Alexander Moschen, Alexander Höller, Wolfgang Schobersberger, Martin Widschwendter

## Abstract

While intermittent fasting (IF) promotes longevity in animal models, its systemic effects in humans remain poorly understood. Here, we present a six-month longitudinal IF intervention in 114 women (BMI 25-35) with deep clinical, molecular, and microbiome profiling across >3,400 biospecimens from six tissues. Analyses spanning >2,200 multi-omic features and 11,000 microbial function predictions demonstrate coordinated clinical benefits, including improvements in body composition and cardiorespiratory fitness, and reveal coordinated molecular responses across tissues. Iron metabolism emerged as a central axis: transferrin increased while ferritin, haemoglobin, and erythrocytes decreased, changes that opposed ageing trajectories yet remained within physiological limits. Epithelial DNA methylation biomarkers (cervical, buccal) of cancer risk reduced, while blood clocks were largely unresponsive, underscoring tissue-specificity of the epigenome. Immune profiling uncovered dynamic, partially reversible shifts. Notably, we derived a new immunophenotyping-based *ImmuneAge* score that increased during fasting and tracked with inflammatory function, while the pro-inflammatory cytokine IL-17A declined selectively in postmenopausal women. Oral microbiota showed rapid restructuring, whereas gut microbiota shifted more subtly toward enhanced metabolic capacity. Together, these data provide unprecedented insight into the systemic and tissue-specific responses to IF in humans and identify iron homeostasis and immune remodelling as candidate mechanisms. Our findings are available through the Lifestyle Atlas (https://eutops.github.io/lifestyle-atlas).

**Highlights:**

- Intermittent fasting remodels iron metabolism, opposing age-associated trajectories
- Epithelial but not blood methylation biomarkers respond to fasting
- Fasting transiently elevates ImmuneAge and TNF-producing cytotoxic T cells
- Oral microbiota restructure rapidly, while gut microbiota show subtler functional shifts
- Integrative networks link iron, immunity, adiposity, and epithelial barrier function

## Introduction

The discovery in the 1930s that caloric restriction could extend lifespan ^1^ inspired the vision that ageing itself might be targeted to delay or prevent the onset of multiple age-related diseases in parallel and thereby extend healthy life years ^2^. Caloric restriction and fasting-mimicking diets, such as intermittent fasting (IF), have since been shown to robustly extend healthspan in multiple animal models ^3^ and to elicit metabolic benefits in humans ^4^, in part via a metabolic switch from glucose to ketone body utilisation. These non-pharmacological interventions that may slow biological ageing are incredibly relevant in the context of a globally ageing population, where they could reduce both individual disease burden and socio-economic costs. However, their long-term effects in humans remain difficult to study: human lifespans are relatively long, individual responses vary, and multi-decade trials are rarely feasible for ethical and logistical reasons.

Quantifiable measures of ageing, posited as early as 1969 ^5^, offer a potential solution. If validated as surrogate endpoints, such biomarkers could enable early evaluation of healthspan-extending interventions ^6^. Several such measures have been proposed. Epigenetic biomarkers based on DNA methylation (DNAme) patterns in blood, buccal, or cervical samples have been shown to predict the risk of cancer ^7–10^, mortality and healthspan ^11–13^, as well as estimate the rate of ageing ^14^. Their adoption could allow both individual- and population-level monitoring of dietary and other geroscience-guided intervention ^6^. Yet, important gaps remain. First, the longitudinal dynamics and responsiveness of these biomarkers to health-promoting interventions, arguably critical for their implementation ^15^, remains poorly defined, with only few trials, such as CALERIE™ ^16^, reporting longitudinal changes. Second, their biological interpretability is limited: it remains unclear which molecular and systemic changes they reflect ^15^, as no study has systematically mapped their associations with other molecular or clinical features during an intervention.

Here, we address these gaps by leveraging a longitudinal, multi-omic dietary intervention study (ClinicalTrials.gov NCT05678426) with matched blood, buccal, cervical, faecal, urine, and saliva samples alongside detailed clinical measurement, wearable data, and immune profiling. This design enabled us to characterise the systemic and tissue-specific effects of IF at unprecedented depth. By integrating clinical, molecular, and microbial layers, we identify coordinated responses across physiology, metabolism, immunity, and the epigenome to six months of IF. More broadly, we provide one of the most comprehensive human resources to date for assessing the responsiveness of healthspan biomarkers to intervention. The resulting Tyrol Lifestyle Atlas ^17^ establishes a benchmark dataset for discovery and cross-study comparison, with immediate value for refining biomarkers and advancing mechanistic understanding of how lifestyle interventions modulate human health.

## Results

### Study design, cohort, and multi-omic profiling strategy

We recruited 114 female participants aged 30-60 years with BMI 25-35 into the IF arm of the TirolGESUND (Table 1). Participants were instructed to follow a 16:8 fasting regimen for six months, restricting food intake to an 8-hour daily window. We focused on female participants due to a particular interest in recently developed epigenetic biomarkers associated with women’s cancers and ageing. To assess whether ketogenic supplementation could amplify the metabolic effects of IF, participants were block-randomised by menopausal status and BMI to receive a daily multi-chain triglyceride (MCT) supplement (K group) or no supplement (I group) (Extended Data Table 1). Primary analyses examined the overall effects of IF, with secondary analyses testing potential synergistic effects of MCT supplementation.

**Table 1.** Participant characteristics by compliance group.

| Characteristic | Overall<br>n = 114 | high<br>compliance<br>n = 66 | medium<br>compliance<br>n = 20 | low<br>compliance<br>n = 10 | dropout<br>n = 18 |
| --- | --- | --- | --- | --- | --- |
| **Age at consent**, Mean (SD) | 46.7 (7.6) | 47.2 (7.4) | 45.3 (9.3) | 49.1 (5.5) | 45.0 (7.5) |
| **BMI at consent** (kg/m<sup>2</sup>), Mean (SD) | 29.7 (3.1) | 29.7 (2.9) | 28.8 (2.7) | 30.6 (3.7) | 30.6 (3.8) |
| Δ M2, Mean (SD) | -0.9 (0.7) | -1.1 (0.7) | -0.7 (0.6) | -0.8 (0.7) | -0.4 (0.6) |
| Δ M4, Mean (SD) | -1.4 (1.1) | -1.7 (1.2) | -1.0 (1.0) | -0.9 (0.6) | -0.7 (0.8) |
| Δ M6, Mean (SD) | -1.7 (1.4) | -2.0 (1.5) | -1.0 (0.9) | -0.8 (0.8) | -- |
| **Smoking history**, n (%) |  |  |  |  |  |
| Never smoker | 65 (57) | 37 (56) | 12 (60) | 4 (40) | 12 (67) |
| Former smoker | 41 (36) | 25 (38) | 6 (30) | 5 (50) | 5 (28) |
| unknown | 8 (7.0) | 4 (6.1) | 2 (10) | 1 (10) | 1 (5.6) |
| **Dietary preference**, n (%) |  |  |  |  |  |
| normal | 100 (88) | 59 (89) | 17 (85) | 9 (90) | 15 (83) |
| vegetarian/vegan/pescetarian | 4 (3.5) | 2 (3.0) | 1 (5.0) | 0 (0) | 1 (5.6) |
| unknown | 10 (8.8) | 5 (7.6) | 2 (10) | 1 (10) | 2 (11) |
| **Weekly intense activity** (min), n (%) |  |  |  |  |  |
| no intense activity | 61 (54) | 38 (58) | 9 (45) | 5 (50) | 9 (50) |
| 30-149 min | 36 (32) | 19 (29) | 7 (35) | 3 (30) | 7 (39) |
| ≥150 min | 9 (7.9) | 5 (7.6) | 2 (10) | 1 (10) | 1 (5.6) |
| unknown | 8 (7.0) | 4 (6.1) | 2 (10) | 1 (10) | 1 (5.6) |
| **Postmenopausal**, n (%) | 37 (32) | 21 (32) | 7 (35) | 4 (40) | 5 (28) |
| **Regular medication use**, n (%) | 19 (17) | 11 (17) | 5 (25) | 2 (20) | 1 (5.6) |
| Antihypertensive medication, n (%) | 11 (9.6) | 8 (12) | 2 (10) | 0 (0) | 1 (5.6) |
| Antidepressant medication, n (%) | 3 (2.6) | 1 (1.5) | 1 (5.0) | 1 (10) | 0 (0) |
| Other medication, n (%) | 6 (5.3) | 3 (4.5) | 2 (10) | 1 (10) | 0 (0) |
| **Systolic blood pressure** (mmHg), Mean (SD) | 116.7 (13.5) | 116.8 (14.2) | 113.5 (12.7) | 120.0 (12.5) | 117.8 (12.4) |
| Δ M6, Mean (SD) | -2.5 (10.7) | -2.7 (10.2) | 0.4 (11.4) | -8.3 (13.3) | -- |
| **Diastolic blood pressure** (mmHg), Mean (SD) | 76.3 (8.3) | 76.4 (8.7) | 75.0 (7.9) | 78.0 (7.5) | 76.9 (7.7) |
| Δ M6, Mean (SD) | -2.4 (7.8) | -1.9 (7.1) | -1.8 (7.3) | -10.0 (12.2) | -- |
| **Total cholesterol** (mg/dL), Mean (SD) | 203.9 (31.9) | 204.1 (31.8) | 195.4 (30.3) | 216.6 (26.8) | 206.0 (36.6) |
| Δ M6, Mean (SD) | 6.9 (21.4) | 6.0 (21.1) | 9.1 (22.7) | 10.2 (23.6) | -- |
| **Triglycerides** (mg/dL), Median (IQR) | 90.0 (51.8) | 91.0 (54.8) | 81.5 (29.8) | 79.5 (38.8) | 108.5 (47.0) |
| Δ M6, Median (IQR) | 6.5 (35.8) | 5.0 (37.0) | 16.0 (44.0) | 9.5 (31.0) | -- |
| **HbA1c** (%), Mean (SD) | 5.3 (0.3) | 5.4 (0.3) | 5.3 (0.2) | 5.3 (0.2) | 5.4 (0.3) |
| Δ M6, Mean (SD) | 0.0 (0.2) | 0.0 (0.2) | 0.1 (0.1) | 0.1 (0.1) | -- |
| **Fasting glucose** (mg/dL), Mean (SD) | 80.5 (13.4) | 79.6 (13.3) | 84.0 (16.7) | 85.6 (12.5) | 76.8 (8.0) |
| Δ M6, Mean (SD) | -0.6 (15.8) | 0.3 (14.2) | -0.9 (20.7) | -7.7 (15.9) | -- |
| **Haemoglobin** (g/dL), Mean (SD) | 13.2 (0.8) | 13.1 (0.8) | 13.2 (0.8) | 13.4 (0.8) | 13.2 (0.7) |
| Δ M6, Mean (SD) | -0.2 (0.6) | -0.2 (0.6) | -0.2 (0.7) | -0.1 (0.4) | -- |

In addition to IF, participants were encouraged to maintain or increase regular physical activity, with optional access to three guided exercise sessions (balance and coordination, strength, and aerobic exercise). Participants self-selected their fasting window, with most choosing to skip breakfast (**Extended Data Figure 1a**). Biological samples from multiple tissues (blood, buccal, cervical, saliva, urine, and faecal) were collected every two months for molecular profiling, alongside routine haemograms and body composition measurements. More detailed clinical assessments, vascular and body fat sonography, and optional dermatological measurements were conducted at baseline and six months (**Figure 1a**). Weekly compliance was recorded by personal coaches and summarised as the percentage of days within each two-month interval and overall (**Figure 1b, c**). Compliance was generally high, though slightly higher earlier in the study, and was categorised as high (fasting for 16 h on ≥85% of days during the entire study), medium (60-84.9%) or low (<60%) (**Figure 1d**; participant characteristics by compliance group and intervention in **Table 1**). Of the 114 participants, 96 (84.2%) completed the study. Dropouts (n=18) tended to show smaller BMI changes at interim follow-ups, but had no other distinguishing baseline characteristics (**Extended Data Figure 1b**, **Table 1**). In the K group, 31/51 individuals (60%) who completed the study reported consistent MCT supplement use, with adherence mirroring fasting compliance (**Extended Data Figure 1c, d**).

**Figure 1.**
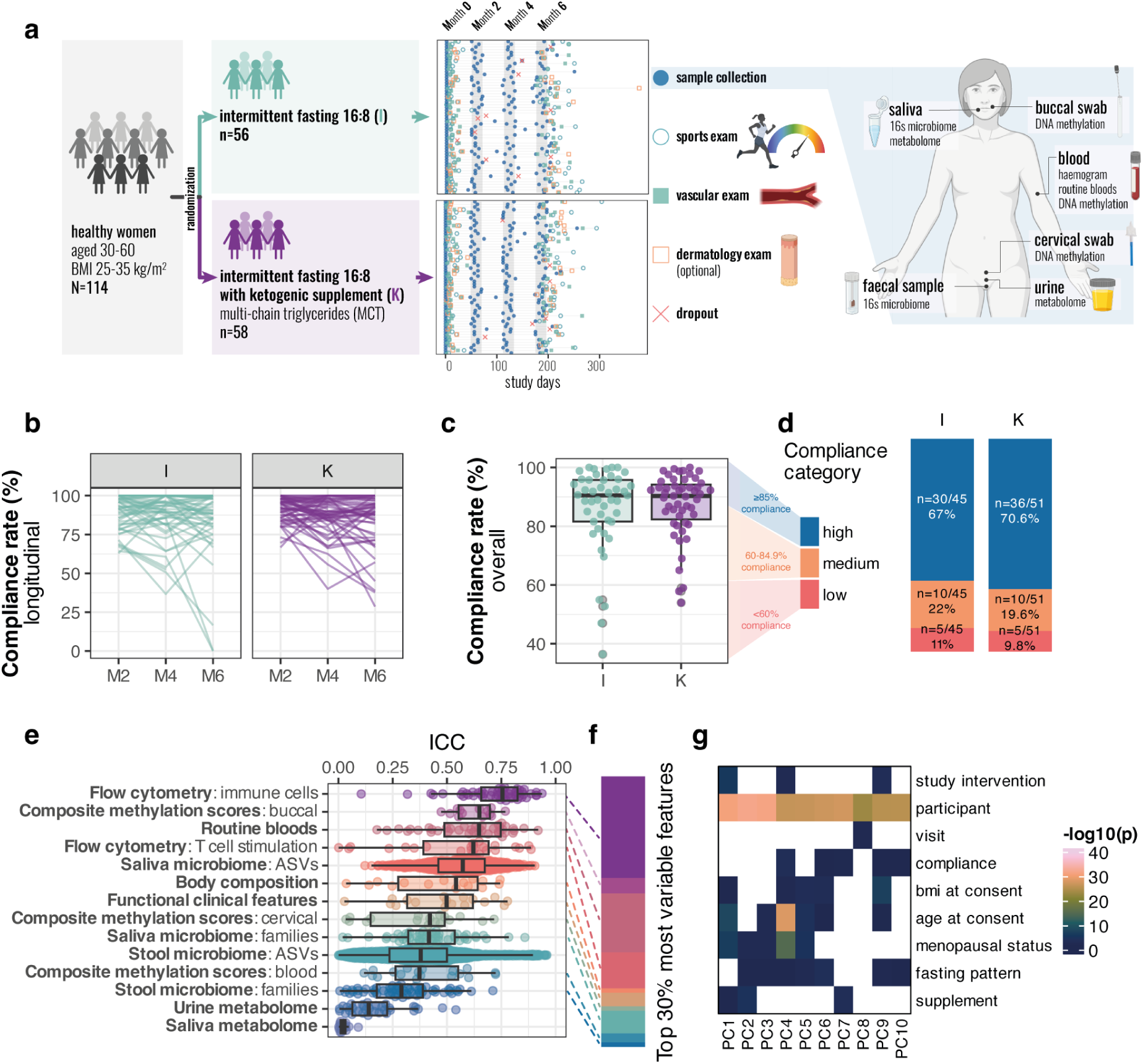
Study overview and compliance. **a** Study design, overview of appointments across individuals by visit (month 0-6), and collected samples. In the appointment diagram, each row represents one individual, and each point a study visit or event. **b** Longitudinal compliance rates of individuals in the two study arms, defined as % of study days that the 16 h of fasting were adhered to in the previous two months. **c** Overall compliance rate across the study in individuals who completed the study. **d** Barplot displays the number of individuals who completed the study in each compliance category (high: ≥85% compliance rate; medium: 60-84.9%; low: <60%). **e** Intraclass correlation coefficient (ICC) of individual features by assay. A high ICC indicates high variability across individuals (i.e., more personal features). **f** Distribution of the assays for the top 30% most personally variable features used for principal component analysis in g (ASVs were excluded). Colour legend is the same as in e. **g** Association of the first five principal components with participant characteristics and study details. P values are coloured by −log10(p), with any p>0.05 shown as blank. **Abbreviations:** I, Intermittent fasting group. K, Intermittent fasting plus multi-chain triglyceride (ketogenic supplement). MCT, multi-chain triglyceride. M0, M2, M4, M6: month 0, 2, 4, 6. ICC, intraclass correlation coefficient. PC, principal component. ASV, amplicon sequence variant. Boxplots indicate median and interquartile range while whiskers indicate values +/− 1.5 interquartile range. Individual data points are shown where possible.

### Intermittent fasting-focused intervention improves perceived health, sleep, and well-being

More than a third of participants reported positive effects of the intervention, such as improved wellbeing, sleep quality, and energy levels (**Extended Data Figure 1e, f**). Negative effects - primarily bloating, nausea, or abdominal discomfort - were more frequently reported in the K group and were predominantly attributed to MCT supplementation. In 56/96 completers with health-related quality of life data at baseline and six months, self-reported health on the EQ-5D-5L ^18^ visual analogue scale (VAS) significantly improved in both I and K groups (p=0.00099 and p=0.018, respectively, paired two-sided Wilcoxon tests), increasing from an average of 80.6 to 88.5 overall (**Extended Data Figure 1g**). As expected in a health cohort, no participants reported self-care limitations at baseline. Across the remaining health-related quality of life domains - activity, anxiety, mobility,and pain - scores improved at six months compared to baseline (**Extended Data Figure 1h**). To assess whether changes might be confounded by shifts in habitual activity, we examined wearable device data in those who completed the study. Average daily steps across two-week windows prior to each visit did not change substantially from baseline (first two weeks per participant) to the intervention period, suggesting that large differences in physical activity were unlikely to explain the observed effects (**Extended Data Figure 1i**).

### Longitudinal multi-omic profiling reveals personal and intervention-driven variability

The collection of longitudinal data allowed us to assess changes over time within and between individuals across several molecular and cellular features. We initially explored the variance explained by the participant structure using intraclass correlation coefficients (ICC) from basic linear mixed-effects (LME) models. Consistent with previous reports ^19,20^, we found that immune-related and clinical features were the most personally distinct, whereas stool microbiome and metabolome profiles showed greater temporal variability (**Figure 1e**). Epigenetic features, such as buccal composite scores, also exhibited high personal stability. Exploring the top 30% of personal features (**Figure 1f**) in more detail, we observed that their first 10 principal components (PCs) captured both inter-individual and intervention-associated variation (**Figure 1g**, **Extended Data Figure 2a-i**).

To systematically disentangle the effects of time, intervention, and compliance, we analysed each data type using both a modified intention-to-treat (all completers) and per-protocol approach stratified by compliance group (**Figure 1d**). For each feature, we assessed change over time using two-sided Wilcoxon tests and linear mixed-effects models adjusting for covariates (**Extended Data Tables 2-8**). We first describe the overall effects of the IF-focused intervention (**Figures 2–5**), then examine potential modulation by ketogenic supplementation, baseline characteristics, and model specific on clinical and molecular features (**Figure 6**). Finally, we integrate the multi-omic data to identify cross-omic associations (**Figure 7**).

**Figure 2.**
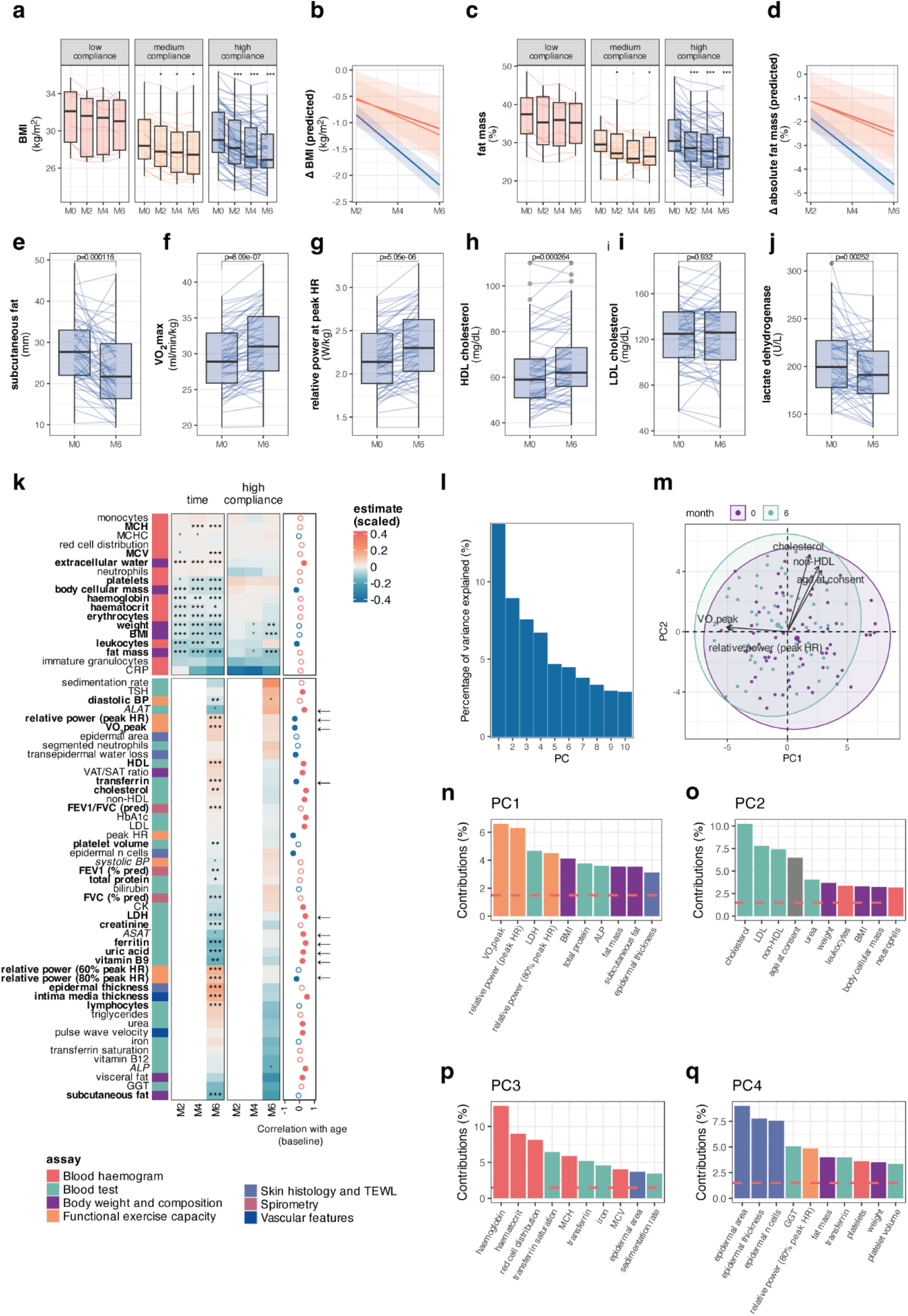
Clinical changes in response to intermittent fasting. **a** Body mass index over time in complete cases of lower, medium, and high compliance groups. p values obtained from paired two-sided Wilcoxon test compared to M0. **b** Predicted average change in body mass index based on time and compliance group, derived from linear mixed effects models adjusted for age at consent and assessing the interaction of time and compliance. **c** Fat mass over time in complete cases of lower, medium, and high compliance groups. p values obtained from paired two-sided Wilcoxon test compared to M0. **b** Predicted average change in fat mass based on time and compliance group, derived from linear mixed effects models adjusted for age at consent and assessing the interaction of time and compliance. **e** Subcutaneous fat, **f** VO_2_peak, **g** relative power at maximum heart rate, **h** HDL cholesterol, **i** LDL cholesterol, **j** lactate dehydrogenase at baseline and month 6 in highly compliant individuals. p values in **e-j** are obtained from paired two-sided Wilcoxon tests and are FDR corrected. **k** Heatmap illustrating estimates for impact of time (value ~ age at consent + bmi at consent + visitId + (1|subjectId)) and interaction between time and high compliance (value ~ age at consent + bmi at consent + visitId*compliance + (1|subjectId)) from linear mixed-effects models. p values were adjusted for multiple testing using both FDR correction and permutation testing. For details see the methods section. P values and corresponding row labels that are both significant at p < 0.05 after FDR correction and permutation testing are shown in bold, whereas p values and row labels that were significant after permutation testing only (but not FDR correction) are shown in grey and italic. *, **, *** indicate p < 0.05, < 0.01, and < 0.001 of uncorrected p values respectively. Right annotation indicates correlation with age at baseline (filled circles are p<0.05). Arrows on the right indicate if the estimate of time (p < 0.05) is going in the opposite direction of the correlation with age at baseline. **l** Scree plot of principal components of clinical data in highly compliant individuals at M0 and M6. **m** Principal components 1 and 2 coloured by time point highlight a general shift over time, associated with VO_2_peak and exercise capacity. Top 10 contributors to **n** PC1, **o** PC2, **p** PC3, and **q** PC4, coloured by assay. Legend for assay colour is the same as in panel k. **Abbreviations**: BMI, body mass index. HDL, high density lipoprotein. LDL, low density lipoprotein. PC, principal component. Boxplots indicate median and interquartile range while whiskers indicate values +/− 1.5 interquartile range. Individual data points are shown where possible.

### Intermittent fasting is associated with pronounced changes in body composition, cardiorespiratory fitness (VO_2_peak), and iron metabolism

Over six months, the 96 participants completing the study lost an average of 4.7 kg (SD 3.9) with greater reductions in BMI and fat mass among those with high fasting compliance (**Figure 2a-d, k**; **Extended Data Tables 2, 3, 6, 7**). Highly compliant individuals reduced their BMI by 6.8% (−2.01 kg/m^2^, SD 1.51) and fat mass by 4.3 percentage points (SD 3.3) (**Extended Data Table 5**), driven largely by subcutaneous rather than visceral fat (**Figure 2e, k; Extended Data Figure 3a, b**). Visceral fat tended to decrease, but changes were small and non-significant, and the ratio of visceral to subcutaneous fat (VAT/SAT ratio), a predictor of cardiometabolic risk ^21^, remained unchanged (**Figure 2k; Extended Data Figure 3c**). Body composition also shifted toward higher extracellular water (+0.67L) and lower body cellular mass (**Figure 2k**; **Extended Data Table 1, 2**), with attenuation in highly compliant participants (**Figure 2k; Extended Data Figure 3d, e**).

Cardiorespiratory fitness improved, with VO_2_peak increasing by 5.6% overall and up to 6.9% in highly compliant individuals (**Figure 2f, k, Extended Data Table 2**). Relative exercise capacity at 60%-100% peak heart rate rose by 7-14% in the highly compliant group (**Figure 2k**, **Extended Data Tables 2, 7**). Serum HDL cholesterol increased, while LDL remained stable (**Figure 2h, i, k**), although neither were modulated by compliance (**Figure 2k**, **Extended Data Figure 3g, h**). Biomarkers of liver and kidney function and cellular damage, including lactate dehydrogenase (LDH) (**Figure 2j**), alanine-aspartate transferase (ALAT), or uric acid (**Figure 2k**; **Extended Data Tables 2, 6**, **Extended Data Figure 3i**) improved, and ALP decreased significantly only in highly compliant participants (**Figure 2k**). C-reactive protein (CRP) tended to fall, and more so in highly compliant individuals, suggesting a potential decrease in inflammation, but this did not reach statistical significance.

IF also impacted iron metabolism: transferrin increased, while ferritin, haemoglobin, haematocrit, and erythrocytes decreased. Values remained within physiological range (**Extended Data Figure 4a-e**), and opposed age-associated patterns of rising ferritin and falling transferrin ^22^, suggesting rejuvenation of iron-related physiology. Effects were independent of meal timing (**Extended Data Figure 4f, g**). Additional effects were related to vitamin B9 metabolism, but likewise remained in physiological ranges (**Extended Data Figure 4h**). Lower systemic iron has been linked to improved longevity in Mendelian randomisation study ^23^, centenarian cohorts showing higher levels of iron binding proteins ^24^, and animal studies where inhibiting iron metabolism extended lifespan ^25^. Blood donation has similarly been associated with reduced mortality (even after adjusting for the healthy donor effect) ^26^ and improved skin ageing via reduced iron deposition ^27^. In line with this, we observed a significant increase in epidermal thickness in skin biopsies (+14.1% from baseline, n=28) without changes in epidermal area or cell number. Evidence on IF and iron is limited, but caloric restriction in mice improved ventricular hypertrophy through increased transferrin receptor and reduced ferritin expression ^28^, consistent with our findings.

Finally, intima-media thickness showed a modest increase, more attenuated in highly compliant participants, without a corresponding change in pulse wave velocity (**Figure 2k; Extended Data Figure 3j, Table 2**), warranting further investigation into vascular effects of fasting.

An overview of the impact of time and compliance effects across clinically relevant features is shown in **Figure 2k** (raw data in **Extended Data Tables 6-8**). An example interaction of fat mass reduction with time, further enhanced by compliance, is illustrated in **Supplementary Figure 1**). PCA of features in **Figure 2k** at baseline and month six across highly compliant individuals revealed four principal components explaining 24.7% of the variance, associated with cardiorespiratory fitness (VO_2_peak, relative capacity at peak heart rate), body composition (body mass index, fat mass, subcutaneous fat), and blood parameters including cholesterol, LDL, lactate dehydrogenase, haemoglobin, platelets, and others (**Figure 2m**). In line with observations from individual variables, PCA highlighted overall improvements in VO_2_peak and BMI (**Figure 2n-q**).

### Intermittent fasting modulates epigenetic biomarkers in epithelial tissues more than in blood

DNA methylation biomarkers have been developed in non-invasive samples, such as blood, cervical, or buccal swabs as indicators for cancer risk (e.g., Women’s cancer risk identification indices, or WID ^7–10,29,30^), age-related diseases (e.g., diabetes ^31^), the rate of ageing (DunedinPACE ^14^), and healthspan or mortality (e.g., PhenoAge ^11^, GrimAge ^12^, CauseAge ^13^). Yet their responsiveness to interventions, and the degree of concordance across tissues, remain less clear. As diet, obesity, and age are known risk factors for several cancers, we investigated the dynamics of epigenetic biomarkers across multiple tissues over six months of IF. Principal components of the top 5% most variable CpGs (Illumina EPIC v1 array) were strongly associated with neutrophil or immune proportions in blood, buccal, and cervical samples, and with visit in the latter two (**Figure 3a**; **Extended Data Figure 5b, e**), confirming expected biological drivers of variation. The first principal component captured most of the variance in buccal and cervical samples (94.2% and 83.7%, respectively), while variance in blood was distributed across more components (**Extended Data Figure 5a, d, h**). Both buccal and cervical samples showed a gradual reduction in immune cell proportions, especially in high-compliance individuals, whereas neutrophil estimates (the major immune cell of whole blood ^32^) remained stable (**Figure 3b, c**; **Extended Data Figure 5b, c, e, i**). These patterns were consistent with reduced immune infiltration in epithelial tissues, in line with reports of reduced periodontal inflammation with fasting ^33^. Downstream analyses therefore adjusted cervical and buccal biomarkers for both immune cell proportion and chronological age.

**Figure 3.**
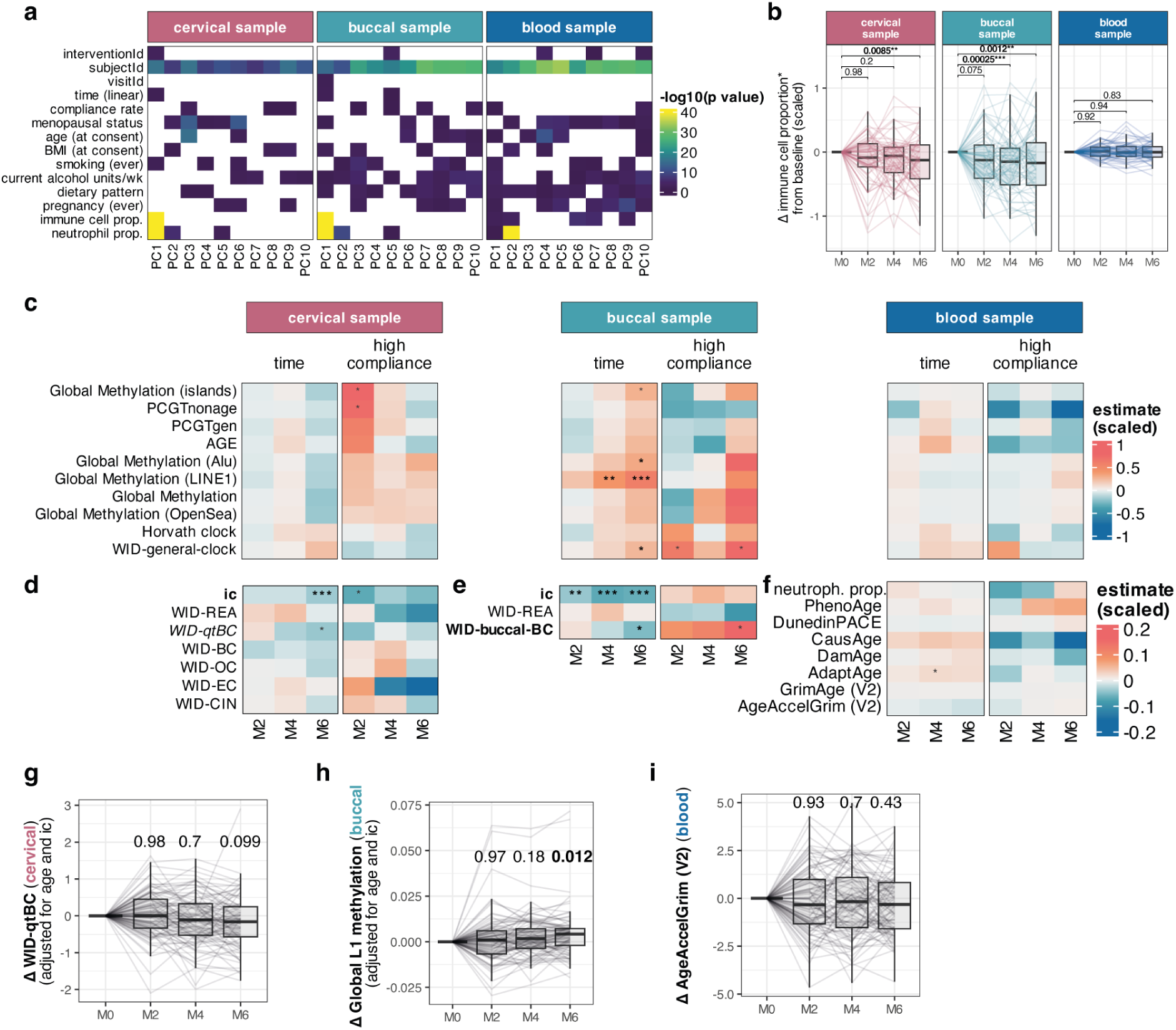
Epigenetic changes in different sample types in response to intermittent fasting. **a** Association of features with principal components highlight variability over time, sample characteristics and phenotypic factors at baseline. Only variables with a significant association (p < 0.05) are coloured (i.e., any cells p > 0.05 are blank). p values are derived from Pearson correlation test (linear covariates) or Kruskall-Wallis test (factorial covariates). Principal components were computed using complete cases only. **b** Change in immune (or neutrophil*) proportion over time. p values derived from paired two-sided Wilcoxon test compared to baseline and FDR corrected. **c** Heatmap illustrating linear mixed-effects model estimates for impact of time (value ~ age at consent + bmi at consent + time + (1|subjectId)) and, interaction between time and high compliance (value ~ age at consent + bmi at consent + time*compliance + (1|subjectId)), on shared DNA methylation biomarkers in different sample types. *, **, *** indicate p < 0.05, < 0.01, and < 0.001, respectively. Bold p values remain significant after FDR correction, while gray p values remain significant after permutation testing. **d** Tissue-specific DNA methylation biomarkers in cervical samples. Scale is the same as in c. **e** Tissue-specific DNA methylation biomarkers in buccal samples. Scale in and e is the same as in c. **f** Tissue-specific DNA methylation biomarkers in blood samples. Bold p values and corresponding row labels in d-f indicate those significant after FDR correction, while grey p values and italic labels are those features that remain significant after permutation testing. **g** WID-qtBC index changes over time in paired cervical samples. p values derived from paired two-sided Wilcoxon test compared to baseline and corrected for FDR. **h i** Global LINE1 (L1) methylation in buccal paired samples over time. p values derived from paired two-sided Wilcoxon test compared to baseline and corrected for FDR. **i** GrimAge (V2) acceleration in paired blood samples over time and corrected for FDR. **Abbreviations**: ic, immune cell proportion. PC, principal component. MCT, multi-chain triglyceride. Boxplots indicate median and interquartile range while whiskers indicate values +/− 1.5 interquartile range. Individual data points are shown where possible.

The WID-qtBC, a breast cancer risk predictor integrating genetic and nongenetic factors ^30^, reduced in cervical samples (**Figure 3d, g**). Other changes were limited, with a transient increase in global methylation and PCGTnonage, a biomarker of stem cell-like methylation ^34^, in high-compliance individuals (**Figure 3d, g**; **Extended Data Figure 5c**). Buccal samples showed increased global methylation at *LINE1* repetitive elements, whose hypomethylation is a biomarker for cancer ^35^ and genome instability ^36^, alongside a reduction in the WID-buccal-BC ^32^ (**Figure 3c, e**), though less pronounced in highly compliant individuals (**Extended Data Figure 5f**). The WID-general-clock also increased, further so in highly compliant individuals (**Figure 3c**; **Extended Data Figure 5g**).

In blood, effect sizes were generally small (**Figure 3f**; note different scale for **Figure 3f**). mITT analysis indicated modest, largely non-significant changes across multiple ageing biomarkers (PhenoAge, AdaptAge, DamAge, CausAge; **Figure 3c, f**, **Extended Data Figure 5j**). DunedinPACE, a rate of ageing biomarker previously reported to slow during two years of caloric restriction ^16^, was not significantly affected by six months of intermittent fasting (**Figure 3f**; **Extended Data Figure 5k**).

Thus, IF primarily modulated epigenetic features in epithelial tissues, including reductions in breast-cancer associated signatures and global methylation, whereas blood biomarkers were largely unresponsive. These findings highlight the tissue specificity of intervention-induced epigenetic change. While the ICC was higher for buccal and cervical sample biomarkers than in blood (**Figure 1f**), these findings are not at odds. Instead, they suggest that biomarkers in epithelial-derived tissues may be more personal and less susceptible to random effects (e.g., noise), and can identify coordinated changes over time more easily.

### Intermittent fasting reversibly increases immune age

Ageing can influence immune function and is associated with a decrease in immune function and plasticity ^37^. Flow cytometry of peripheral blood revealed rapid and dynamic shifts in immune cell populations. In line with high intraclass coefficients (**Figure 1e**), PCA showed subject-specific immune profiles but also pronounced changes within two months, with a gradual return toward baseline thereafter (**Extended Data Figure 6a-c**), indicative of transient intervention effects. Several populations changed transiently (e.g., CD94^+^CD4^+^ T cells, B cells, defective NK cells, CD8^+^ T cells), whereas others shifted more gradually over six months (e.g., non-classical monocytes, CD57^+^KLRG1^-^CD8^+^ T cells, CD8^+^ central memory (T_cm_) cells). As expected, many immune populations correlated with age at baseline, and some shifted in the opposite direction after fasting. For instance, CD57^+^KLRG1^-^CD8^+^ T cells increased with age but were reduced after 4 and 6 months, while CD57^-^KLRG1^-^CD8^+^ T cells decreased both with age and intervention (**Extended Data Figure 6e**).

To summarise these alterations, we derived an ImmuneAge score using penalised elastic net regression on baseline data of cell proportions. This metric consisted of 16 cell populations (**Figure 4a**), correlated with chronological age at baseline (training data) and follow-ups (validation); (R=0.62 at baseline, p<0.001, **Figure 4b**) and, unexpectedly, increased significantly during fasting (**Figure 4c**). In individuals with low or medium compliance, ImmuneAge values returned to baseline by month 6 (**Figure 4b**), suggesting reversibility. Functional cytokine assays revealed additional changes and supported this interpretation: TNF production increased in both stimulated and unstimulated cytotoxic T cells, consistent with its known rise in ageing in industrialised populations ^38^, and the frequency of TNF-producting cytotoxic T cells correlated strongly with ImmuneAge (**Extended Data Figure 7a-d**). This suggests that ImmuneAge, although derived on chronological age alone, and leveraging primarily immune count data, captured not only age-associated cellular composition but also pro-inflammatory aspects of immune function. In contrast, IL-17A production rose at month 2 in unstimulated cells but decreased after stimulation, and was not correlated with ImmuneAge (**Extended Data Fig. 7c, e**), indicating that ImmuneAge reflects some but not all functional pathways.

**Figure 4.**
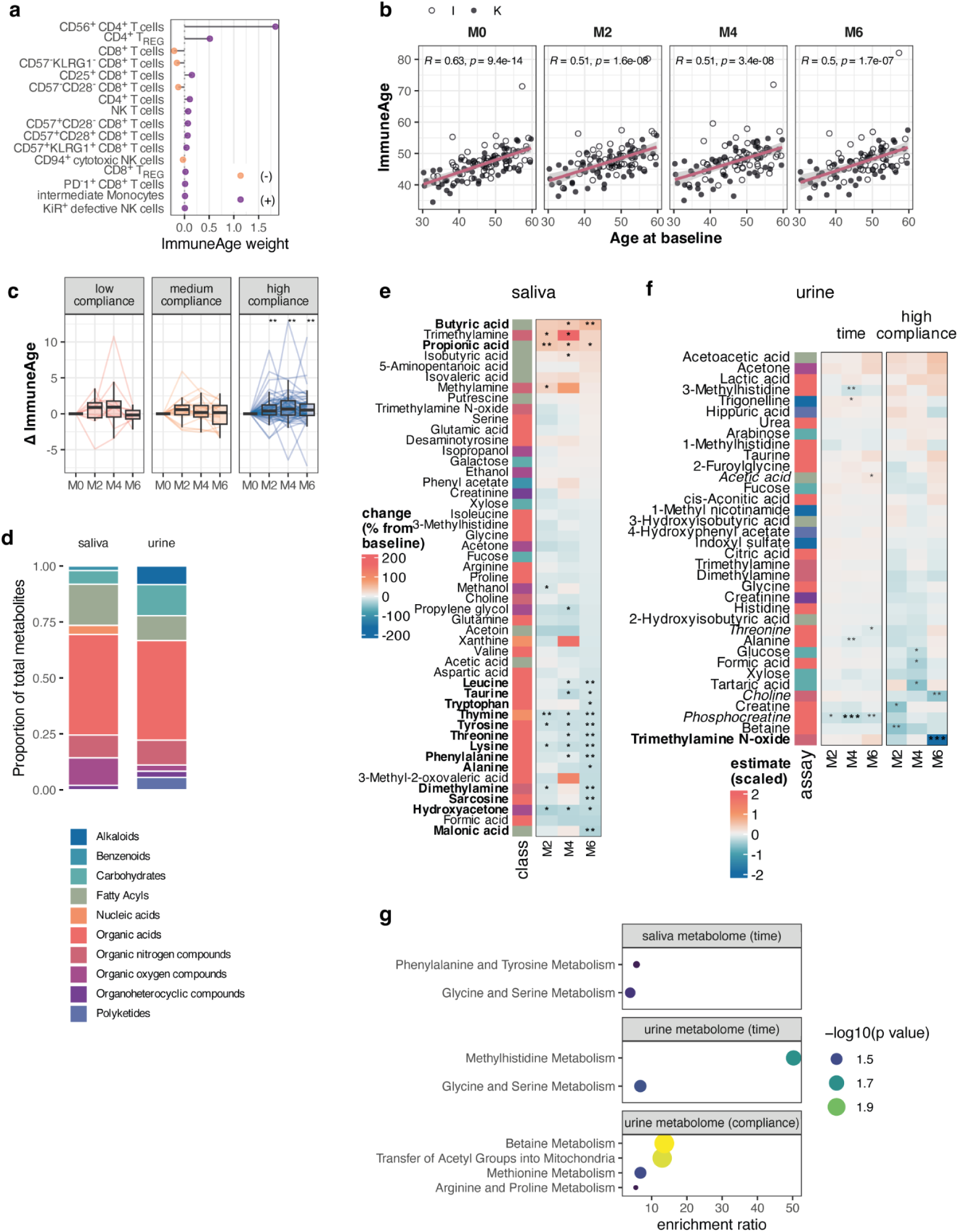
Immune age and metabolomic changes during intermittent fasting. **a** Immune cell populations contributing to composite ImmuneAge. **b** ImmuneAge association with age at baseline. R value denotes the Pearson correlation coefficient. **c** Δ ImmuneAge from baseline by compliance groups. p values are derived from paired two-sided Wilcoxon tests compared to baseline after FDR correction. **, p<0.01; ***, p<0.001. **d** Overview of metabolites identified in saliva and urine samples. **e** Heatmap of saliva metabolites significantly changing from baseline over time (paired two-sided Wilcoxon test compared to baseline, FDR corrected). **f** Heatmap of urine metabolites significantly changing from baseline over time and their modulation by compliance (linear mixed-effects models; FDR correction and permutation testing). P values that remained significant after FDR correction are shown in bold, while p values that remained significant after permutation testing are shown in grey. Features with significant permutation testing-corrected changes at M6 are indicated with italic labels. Annotation of metabolite classes is the same as in e. *, **, *** denote p<0.05, <0.01, and <0.001, respectively. **g** MetaboAnalystR pathway enrichment analysis of significant features in e and f. **Abbreviations:** I, intermittent fasting. K, intermittent fasting plus ketogenic supplement. Boxplots indicate median and interquartile range while whiskers indicate values +/− 1.5 interquartile range. Individual data points are shown where possible.

These data indicate that IF transiently shifts the immune system towards a more pro-inflammatory profile, with both compositional and functional signatures. ImmuneAge emerges as a potential integrative biomarker of immune ageing and function, though immune responses to the IF regime were heterogeneous across cell types and cytokines.

### Metabolic shifts in saliva and urine highlight amino acid catabolism, one-carbon metabolism, and ketogenesis

Untargeted nuclear magnetic resonance (NMR) metabolomics identified 49 metabolites in saliva and 36 in urine, including fatty acids, carbohydrates, organic acids, organic nitrogen compounds (**Figure 4d**). Variance decomposition revealed a low ICC of the saliva metabolome (**Figure 1e**), consistent with influence from recent food intake, whereas the urine metabolome showed modest within-person stability (**Extended Data Figure 8a-c**).

In saliva, 20/49 metabolites (40.1%) changed significantly over time after FDR correction (**Figure 4e**). Short-chain fatty acids (SCFAs; butyrate, propionate, isobutyrate, 5-aminopentanoic acid) increased, likely reflecting oral microbiome fermentation products, while malonic acid decreased. SCFAs in saliva may be linked to both adverse oral health (e.g., tooth decay) and beneficial systemic effects, as observed for gut-derived SCFAs ^39,40^. Nine of 22 organic acids decreased, including several essential (leucine, tryptophan, threonine, lysine, phenylalanine) and non-essential amino acids (tyrosine, alanine, sarcosine), as well as taurine, consistent with reduced intake or altered absorption during fasting.

In urine, 6/36 metabolites (16.6%) changed overall, and 12/36 (33.3%) were modulated by higher compliance (**Figure 4f**). Organic acids, such as 3-methylhistidine (a marker of white meat intake and muscle turnover ^41^), threonine, alanine, and phosphocreatine were reduced, while acetate increased, consistent with enhanced ketogenesis. Compliance-modulated metabolites included reductions in glucose, formic acid, and tartaric acid at month 2, choline, creatine, betaine, and trimethyl-amine N-oxide. These reductions in urine glucose and betaine, and increases in acetic acid were consistent with improved glucose handling, altered one-carbon metabolism, and enhanced ketogenesis.

Pathway enrichment analysis supported these findings (**Figure 4g**). Saliva changes were enriched for phenylalanine/tyrosine and glycine/serine pathways, while urine changes implicated methylhistidine and glycine/serine metabolism. Higher compliance was further associated with betaine and methionine metabolism (one-carbon pathways) and arginine/proline metabolism (**Figure 4g**).

### Rapid restructuring of the oral microbiome and subtler functional shifts in the gut with intermittent fasting

We profiled saliva and stool microbiomes using 16S rRNA gene sequencing, focusing primarily on family-level taxonomic information of the obtained amplicon sequence variants (ASVs) (i.e., the lowest level confidently assigned to most ASVs). PCA showed that both families and ASVs in saliva and faecal samples were strongly personal, with age and menopausal status also contributing to baseline variance (**Figure 5a, b, Extend Data Figure 9a, b**). Lifestyle factors, including previous pregnancies, smoking behaviour, night shifts, alcohol consumption, and fasting pattern (dinner/breakfast skipping) were additionally associated in particular with the oral community structure.

**Figure 5.**
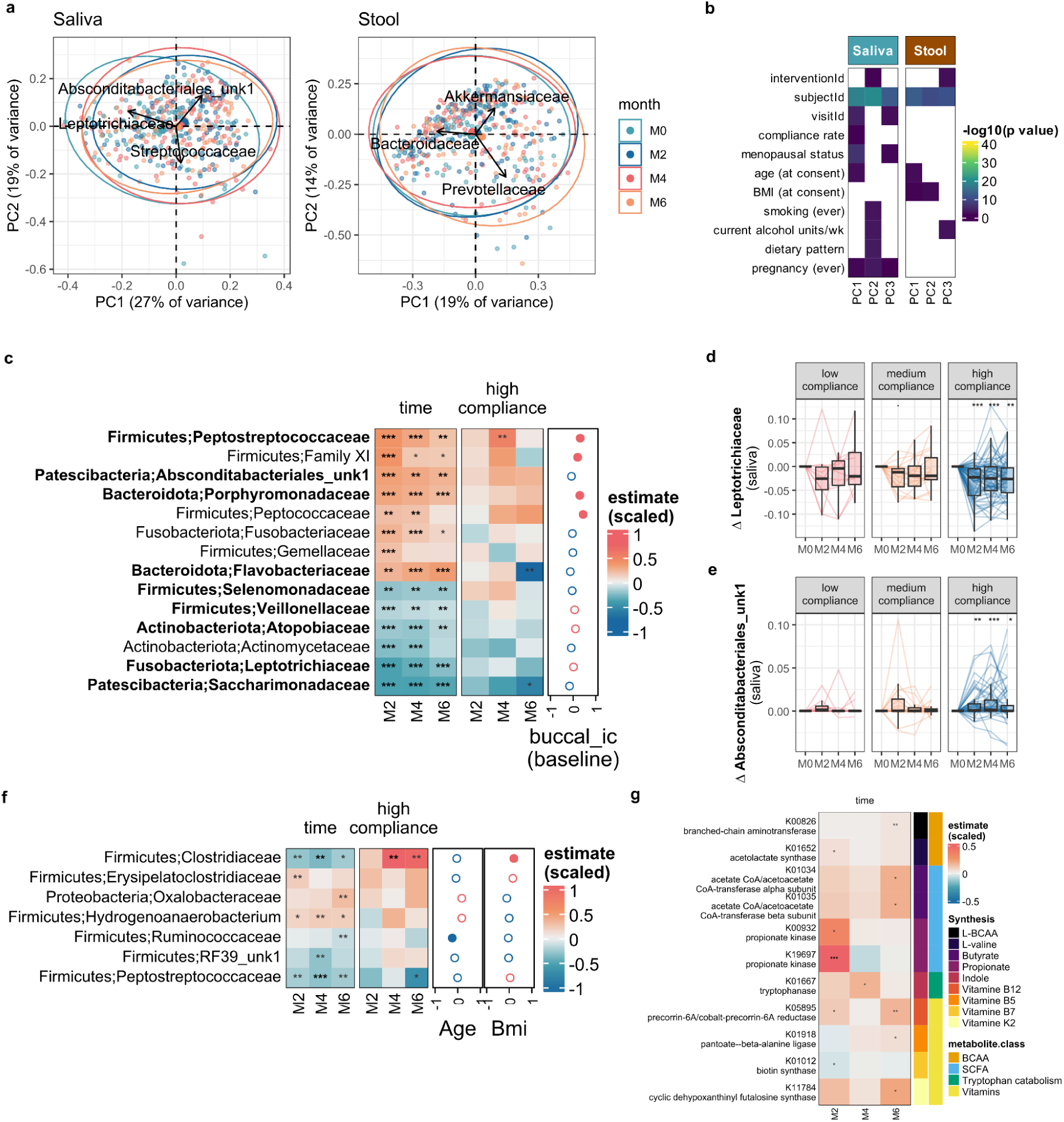
Oral and gut microbiome changes in response to intermittent fasting. **a** Biplots summarise the principal component analyses done on Hellinger transformed count data collapsed at the microbial family-level for the saliva and stool sample sets. Arrows depict the strength and direction of the top three features contributing to the overall variance. **b** Heatmap showing the significance of the first three principal components with recorded covariates. P values > 0.05 appear blank. **c** Significant time (p< 0.01) and corresponding time:high compliance interaction estimates for the differentially abundant oral microbial families compared to baseline from a minimal linear mixed effect model ((value ~ age at consent + bmi at consent + time + (1|subjectId)) and more complex model including a compliance term (value ~ age at consent + bmi at consent + time*compliance + (1|subjectId)),respectively. Pearson correlation of predicted ic content from buccal epigenomes with relative abundance at baseline for the respective families is depicted on the right (full circles indicate p<0.05). **d-e**. Example box plots showing the changes from baseline for selected features. p values are derived from paired two-sided Wilcoxon. **f** Significant time (p value < 0.01) and corresponding time:high compliance estimates for the differentially abundant stool microbial families compared to baseline from a minimal linear mixed effect model and more complex model including a compliance term, respectively. Pearson correlation of age and bmi at consent with relative abundance at baseline for the respective families is depicted on the right (full circles indicate p< 0.05). **f** Time estimates for the differentially abundant predicted gut enzymes (p<0.01) that are implicated in branched chain amino acid metabolism (BCAA), short chain fatty acid production (SCFA), tryptophan catabolism and vitamin metabolismin high compliance individuals. **c**-**g** Stars indicate FDR corrected p values with *, **, *** p<0.05, <0.01, and <0.001, respectively. **Abbreviations:** I, intermittent fasting. K, intermittent fasting plus ketogenic supplement. MCT, multi-chain triglycerides.

Alpha diversity was stable overall, though Chao1 richness tended to decrease in saliva and Shannon diversity to increase in stool (**Extended Data Figure 10a, b**). No changes were observed in the Firmicutes/Bacteroidota and Clostridia/Bacteroidia ratios - two commonly evaluated gut microbiome indicators in the context of obesity and gastrointestinal disorders ^42,43^ (**Extended Data Figure 9c**). Because lower oral microbiome diversity is generally viewed favourably while the opposite is true for the gut ^44^, observed trends in diversity hinted at modest beneficial effects of IF on microbial diversity across tissues. Salivary richness also correlated with buccal cell immune infiltration as estimated from the buccal methylome (r=0.42, p<0.001), supporting links between higher oral diversity, poorer oral health, and inflammation (**Extended Data Figure 10d**).

Differential abundance analysis revealed rapid and sustained changes in the oral microbiome within 2 months of IF **(Figure 5c**). Leptotrichiaceae decreased, while SR1 (an uncharacterised family from the order of Absconditabacteriales) and Streptococcaceae increased (**Figure 5a, d, e**). These effects were particularly pronounced in highly compliant individuals, demonstrating their variability was primarily driven by IF. Porphyrmonadaceae and Veillonellaceae also shifted: *Porphyromonas pasteri*, which increased with increasing compliance, has been associated with periodontal health ^45,46^, whereas *Veillonella* species, decreasing with fasting, are considered accessory pathogens supporting biofilm maturation ^47,48^.

By contrast, stool microbiome changes were slower and smaller (**Figure 5b**, **Extended Data Figure 9f**), and menopause, age, and bmi at consent (baseline) were more contributors to variance than time or intervention group (**Figure 5b**). Families such as Bacteroidaceae, Prevotellaceae and Akkermansiaceae, which constituted the top contributors to PC1 and PC2, were not differentially abundant over time (**Figure 5b**). However, two families that were positively associated with BMI at baseline (Clostridiaceae and Peptostreptococcaceae) decreased over time (**Figure 5f**, FDR corrected p<0.05 at M4). Akkermansiaceae, largely constituted from the emerging probiotic *Akkermansia muciniphila*, a mucin-degrading commensal linked to gut barrier function ^49^, homeostatic immunity ^50,51^ and negatively associated with BMI in humans ^52^, showed substantial inter-individual variability but was not significantly modulated by fasting. However, functional predictions suggested a modest increase in gut microbial capacity for SCFA, branched-chain amino acid, and vitamin biosynthesis in highly compliant individuals (**Figure 5g**). In particular, predicted butyrate metabolism, with known beneficial effects on lipid and glucose homeostasis, appetite regulation, and gut barrier integrity ^40,53,54^, increased in highly compliant individuals.

Together, these findings indicate that the oral microbiome undergoes rapid restructuring during IF, including shifts in taxa linked to oral health, while the gut microbiome shows slower compositional changes but potential functional adaptations that may support metabolic health.

### MCT supplementation selectively alters epigenetic, urine metabolome, and microbial features

Participants were randomly allocated to receive a ketogenic supplement (MCT) or not (**Figure 1a**, **Extended Data Table 1**). The impact of supplementation in addition to IF alone varied considerably across biological systems, with the most affected layers at month 6 being methylation in cervical samples, urine metabolome, and the microbiome (**Figure 6a**). Supplementation did not alter BMI or weight loss, though the loss of fat mass appeared delayed in supplemented individuals (**Figure 6b**, **Extended Data Table 4**). Other clinical parameters, including platelets, diastolic blood pressure, and vitamin B12, were also modestly modulated.

**Figure 6.**
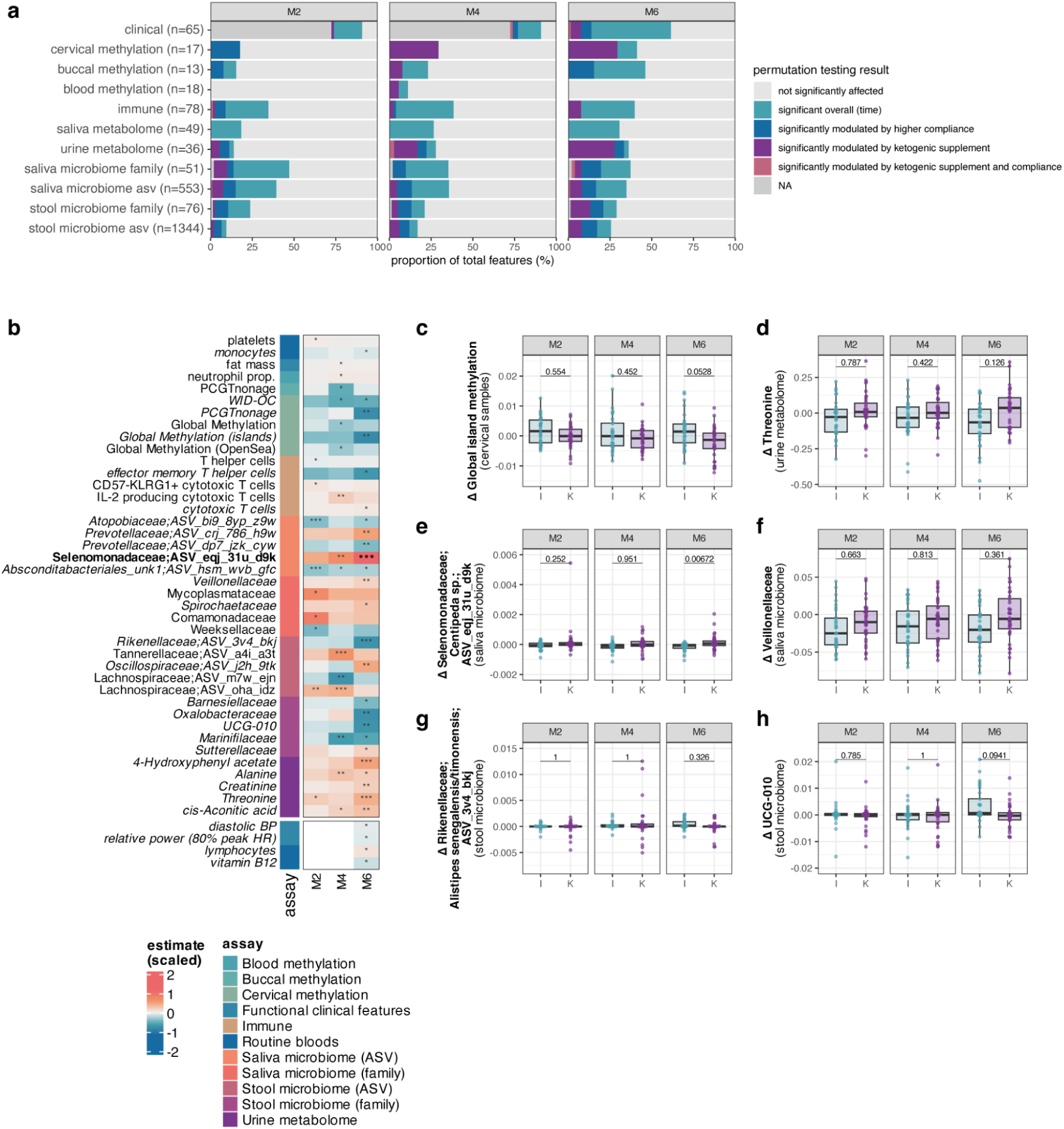
Modulation of biological features by ketogenic supplementation. **a** Overview of the proportion of features across different feature groups that were significantly altered by time and/or modulated by higher compliance or ketogenic supplementation after FDR correction (saliva metabolome; Wilcoxon tests) or permutation correction (all other groups; linear mixed-effects models). The number in brackets indicates the total number of features in each group. **b** Heatmap of the modulation estimate [value ~ age at consent + bmi at consent + time*interventionId + (1|subjectId)] of ketogenic supplementation in the top features within each feature group for the highly compliant participants only. p values in bold remain significant after FDR correction. All other p values remained significant after permutation correction. Labels in bold indicate features with significant FDR corrected changes at M6, while labels in italic indicate features with significant permutation-corrected changes at M6. **c-h** Example boxplots showing the changes from baseline in I (intermittent fasting) versus K (intermittent fasting with ketogenic supplementation) groups for selected features. p values shown are obtained from two-sided Wilcoxon tests (FDR adjusted within each “ome”).

In cervical samples, ketogenic supplementation consistently reduced methylation biomarker values relative to the unsupplemented group (**Figure 6b**). For example, average global island methylation increased in the unsupplemented group but decreased with supplementation, leading to a near-significant group difference after 6 months (FDR=0.05; **Figure 6c**). In the urine metabolome, several metabolites were modulated by supplementation (**Figure 6b, d**). Threonine decreased in the unsupplemented group but increased with supplementation, with additional changes in 4-hydroxyphenylacetate, alanine, creatinine, and cis-aconitic acid, all increasing in the supplemented group relative to the unsupplemented group. In the microbiome, MCT supplementation modulated both oral and gut communities, however the only significant time x MCT interaction at month 6 (FDR<0.05) was an increased abundance of *Centipeda* sp. in saliva of participants in the supplemented group (**Figure 6e**). Supplementation also tempered fasting-induced compositional shifts: reductions in Veillonellaceae in saliva were less pronounced with MCT, while increases in *Alistipes* spp., Oxalobacteraceae and Clostridia UCG-010 in stool were only altered in the unsupplemented group (**Figure 6f-h**).

Together, these results indicate that ketogenic supplementation did not affect weight loss but selectively modulated cervical methylation trajectories, urine metabolites, and microbiome composition, sometimes attenuating fasting-induced shifts.

### Menopause shapes cervical epigenetic profiles and modulates immune, microbial, and metabolite responses to fasting

Menopause exerts wide-ranging effects on biological systems and was a prominent driver in principal component analysis of multiple omic layers. We therefore separately examined whether menopausal status modulated responses to IF in highly compliant participants using linear mixed-effects models.

At baseline and across timepoints, several cervical DNA methylation biomarkers were associated with menopausal status after accounting for study group, age, and BMI at consent (**Extended Data Figure 11a; Extended Data Table 9**). The WID-EC index ^9^, a biomarker of endometrial cancer (which rises postmenopause), remained significantly elevated in postmenopausal compared to premenopausal after FDR correction (**Extended Data Figure 11b**). The WID-relative-epithelial-age (WID-REA) ^29^ also tended to be higher in postmenopausal women at most timepoints, though this did not remain significant after multiple testing correction (**Extended Data Figure 11a, c**). These findings confirmed menopause is a prominent factor impacting cervical DNA methylation even after accounting for chronological age. Menopause also modulated responses to IF, though none of these effects survived FDR correction. Buccal immune cell proportions decreased significantly in premenopausal but not postmenopausal women, while IL-17A production in stimulated T helper cells showed the opposite pattern, declining in postmenopausal but not premenopausal women (**Extended Data Figure 11d, e**). IL-17A has been implicated in bone loss in postmenopausal women, suggesting that its reduction during fasting could reflect selective dampening of inflammatory pathways relevant to bone health ^55^. The oral microbiome also showed differential responses, with Saccharimonadaceae (associated with periodontal disease ^56^) decreasing mainly in premenopausal women (**Extended Data Figure 11f**). In the gut, Izemoplasmatales_unk1 and Butyricicoccaceae diverged, with fasting increasing Butyricicoccaceae in premenopausal but not postmenopausal women (**Extended Data Figure 11g**). Postmenopausal women also exhibited lower urinary acetic acid (**Extended Data Figure 11a**).

Together, these data indicate that menopause shapes both baseline cervical epigenetic profiles and may influence dynamic immune, microbial, and metabolite responses to IF, underscoring the need to stratify by menopausal status when evaluating fasting responses, and highlighting the importance of early prevention and tailored interventions across age groups.

### Molecular and clinical changes occur independent of BMI change

Although our study was not designed to specifically assess weight loss, participants lost an average of 4.7 kg (**Table 1**, **Figure 2a**). We therefore tested whether change in BMI (Δ BMI, month 0-6) was associated with other features, using models additionally accounting for baseline age, baseline BMI, and the interaction between compliance and visit. After FDR correction only BMI and fat mass remained significant, indicating that Δ BMI had little independent effect on the large majority of features beyond what was already captured by other covariates.

### Baseline defective NK cells predict BMI change but not dropout

Individuals with higher compliance tended to have slightly lower baseline BMI (**Table 1**, **Figure 2a**), suggesting that those with poorer health at baseline may be less likely to adhere, though potentially gaining more benefit from fasting. Baseline BMI was significantly associated with subsequent reduction in BMI after adjusting for age, intervention, and compliance (p=0.015). After FDR correction, the only baseline feature significantly associated with weight loss was the proportion of defective NK cells, with higher levels predicting greater BMI reduction (**Extended Data Figure 12, Extended Data Table 11**).

We also examined predictors of study dropout. Individuals who discontinued the study tended to show smaller weight loss (**Extended Data Figure 1b, Table 1**), but logistic regression incorporating baseline features or early changes at month 2 revealed no significant predictors after FDR correction, indicating that dropout was not systematically driven by baseline characteristics or early responses.

### Results are consistent across categorical and continuous time models

Our primary analyses modelled time as a categorical variable to capture non-linear dynamics and facilitate visualisation. Sensitivity analysis using ordinal or continuous representations of time yielded highly consistent results, with similar features passing FDR correction and nearly identical effect estimates (e.g., R^2^ of linear time estimate (ordinal) and estimate of visit M6 = 0.97, p<2.2^-16^; **Extended Data Figure 13**). These findings indicate that our results are robust to the choice of time parametrisation.

### Cross-system networks link microbial, metabolic, immune, and clinical changes during fasting

To capture systemic effects of the intervention, we next aimed to integrate molecular and clinical features across time. As an overview, we first visualised the timing of significant changes, revealing early responses (e.g., body composition) versus slower dynamics (e.g., cervical methylation) (**Figure 7a**). To formally integrate multi-omics dynamics, we applied MEFISTO (Method for the Functional Integration of Spatial and Temporal Omics data) ^57^, which identified 15 latent factors in our dataset. Several factors were associated with time (Factors 9, 11, 12, and 15), and Factor 12 was also associated with compliance and study group allocation (**Extended Data Figure 14a, b**). Factor 12 linked saliva microbiota families, buccal immune cell infiltration, and red blood cell parameters (haemoglobin, erythrocytes) (**Extended Data Figure 14c, d**), consistent with our earlier findings implicating microbiota, immune, and iron alterations (**Figure 2k**, **Figure 5c, f, Extended Data Figure 11d**). Factors 9 and 11 were dominated by saliva microbiota and metabolites, including SCFAs such as butyrate, propionate, and acetate, highlighting oral health as a key dynamic component of fasting responses (**Extended Data Figure 14e-h**).

**Figure 7.**
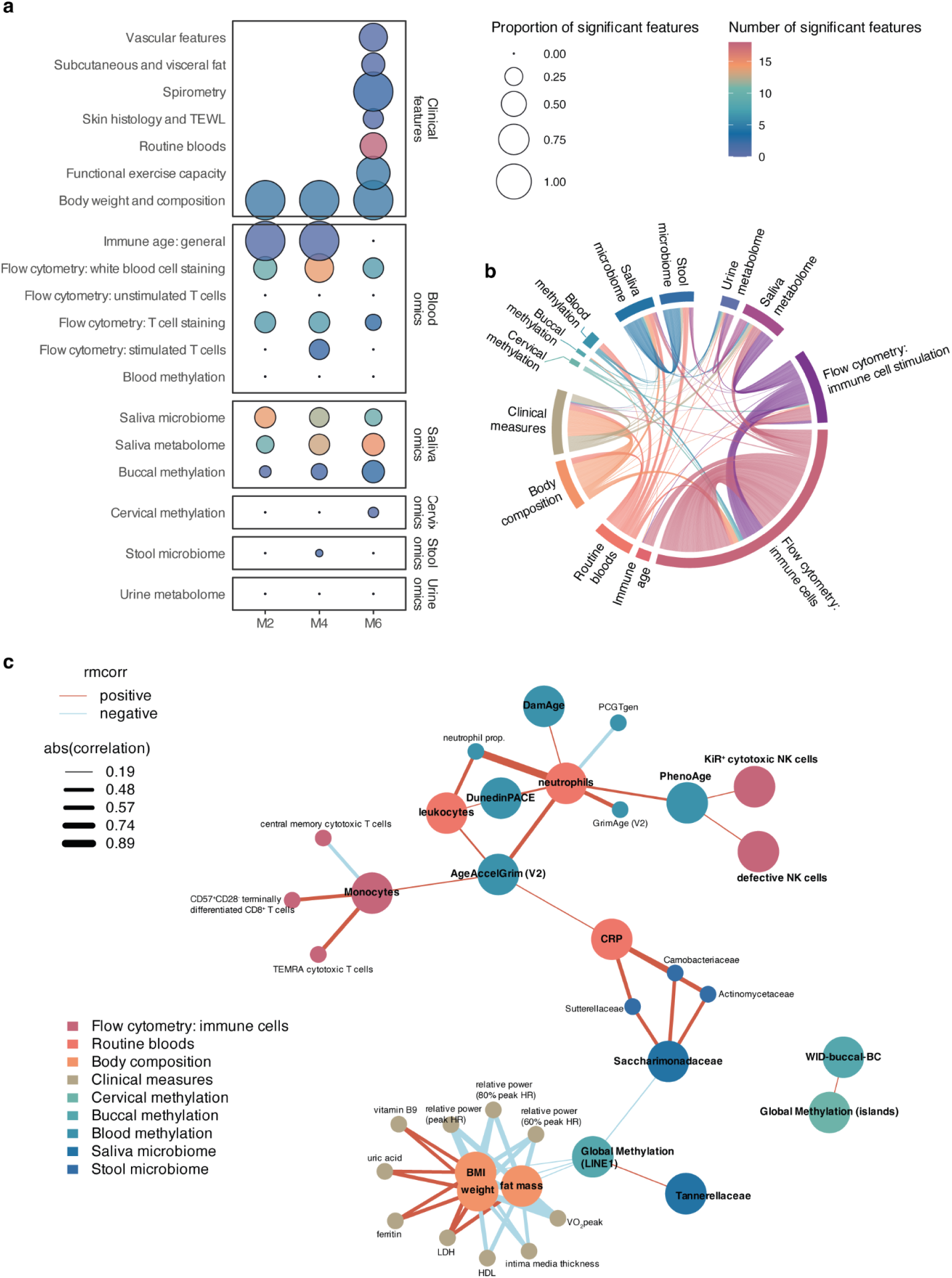
Integration of multi-omic and clinical features identifies correlated changes across multiple systems. **a** Overview diagram visualising timing and number of changes across different omic and clinical features at FDR corrected p<0.05 in linear mixed-effects models (time estimate) or paired two-sided Wilcoxon tests (metabolome data only). **b** Chord diagram visualising cross-omic correlations derived from repeated measures correlation analysis, including correlations at FDR corrected p<0.05 for each feature. **c** Correlation network of selected epigenetic biomarkers and their multi-omic correlates. Nodes shown in bold are the seed features (i.e., epigenetic biomarkers) and their directly connected measures included during expansion (|r|≥0.45), whereas features not in bold include those one step removed. **Abbreviations**: TEWL, transepidermal water loss. r_rm_, repeated measures correlation.

We next applied repeated measures correlation to identify longitudinal associations across omic layers (**Figure 7b, Extended Data Figure 15a, Extended Data Table 12**). This approach confirmed known relationships such as the correlation between methylation-based and measured blood cell counts and between body cell mass and creatinine ^58^ (**Extended Data Figure 15b**, **c**). It also recapitulated expected associations of blood-based epigenetic biomarkers with neutrophils, monocytes, and CRP (**Figure 7c**, **Extended Data Fig 15d-f, Extended Data Table 12**). Conversely, it revealed that buccal and cervical biomarkers did not show the same associations, and for instance global buccal methylation at LINE1 elements associated with the salivary microbiome as well as body composition metrics (**Figure 7c**). Beyond these expected patterns, the analysis revealed key multi-omic hubs with broad cross-system connectivity. Fat mass and ImmuneAge exhibited the largest number of significant correlations. Fat mass was associated not only with other body composition features but also with immune populations, clinical parameters, and microbial families. ImmuneAge correlated with immune populations as expected, but also with subcutaneous fat and specific saliva microbiome families, suggesting coordination between adiposity and immune ageing trajectories. Salivary Saccharimonadaceae, which were found to correlate with buccal global methylation at LINE1 element (**Figure 7c**), emerged as another hub, correlating with CRP, haemoglobin, haematocrit, and epidermal thickness, pointing to an axis connecting oral microbiota, inflammation, iron homeostasis, and epithelial integrity. Other notable associations included a negative correlation between Atopobiaceae and HbA1c, consistent with reduced abundance of this family in diabetes ^59,60^ (**Extended Data Table 12**).

Given the prominence of iron-related changes appeared across several analyses, we specifically explored iron-associated correlations (**Extended Data Figure 16**). Iron parameters were strongly connected with VO_2_peak, BMI, creatinine, uric acid, and LDH, as well as microbial families. Notably, transferrin correlated with pro-inflammatory TNF-producing cytotoxic T cells, reinforcing links between iron handling and immune ageing. These findings aligned with MEFISTO factor 12 and further support the interplay between iron metabolism, immune responses, and systemic physiology.

These integrative analyses highlight cross-omic pathways through which IF exerts systemic effects, converging on interconnected axes of iron metabolism, immune ageing, adiposity, oral microbiota, and epigenetic change. These networks provide a framework for interpreting intervention responses and identifying candidate pathways for future mechanistic studies.

## Discussion

This study provides the most comprehensive multi-omic investigations of IF in healthy women to date, combining clinical, molecular, and microbial profiling across multiple tissues of the same individuals. By repeatedly sampling the same participants, we capture a multi-layered view of how fasting and ketogenic supplementation modulate physiology and age-related processes. This not only enabled identification of changes across systems evolving on different timescales, but also revealed systemic axes linking diverse tissues. The design also allowed comparison with a parallel smoking cessation study ^61^, providing a model for studying diverse health behaviours through harmonised multi-omic profiling.

Over six months, IF produced coordinated improvements in body composition, cardiorespiratory fitness, and favourable shifts in liver and kidney biomarkers (**Figure 2**). Strikingly, we observed consistent alterations in the iron metabolism - an increase in transferrin with decreases in ferritin, haemoglobin, and erythrocytes - opposing age-associated trajectories with remaining within physiological range (**Figure 2k, Extended Data Figure 4**). Integrative analyses reinforced iron biology as a systemic hub, connecting immune activation, VO_2_peak, BMI, and microbial taxa (**Extended Data Figure 14-17**). These findings converge with prior evidence linking reduced systemic iron to improved longevity ^22,23,25,27,28^, and suggests that iron homeostasis is a cross-cutting axis of metabolic, immune, and epithelial barrier function that represents a candidate mechanism of fasting-associated health benefits,

Epigenetic biomarkers responded in a tissue-specific manner. Cervical and buccal samples showed reductions in breast-cancer associated risk signatures and increases in *LINE1* methylation (**Figure 3**), while blood-based clocks, including DunedinPACE - reported to slow after two years of caloric restriction ^16^ - remained largely unchanged. These findings underscore the importance of tissue context: buccal and cervical samples may act as sentinels of dietary and inflammatory exposures, whereas blood may be less responsive over short timeframes. More broadly, they highlight the challenge of applying current epigenetic biomarkers to intervention trials: many clocks trained on cross-sectional cohorts can be insensitive to short-term change, and technical noise can obscure subtle effects. While computational suggestions to bolster technical precision have been proposed ^62^, our prior findings highlight they have their own trade offs, including reduced accuracy ^63^. Refinement of biomarker models through identification of features that are i) reflective of underlying ageing, including in longitudinal data, ii) predict disease outcomes, and yet iii) are malleable through targeted interventions, in combination with minimising technical noise, will be essential for epigenetic biomarkers to become powerful surrogate endpoints in intervention studies ^6,64^.

Immune profiling revealed rapid and partially reversible changes. Flow cytometry showed transient shifts in several T cell and NK subsets (**Figure 4**). An ImmuneAge score, which captured 16 immune populations and was derived in this study, unexpectedly increased during fasting. This coincided with an increase in TNF-producing cytotoxic T cells, consistent with fasting engaging both compositional and functional arms of the immune system (**Figure 4**, **Extended Data Figures 7, 6**). While chronic caloric restriction reduces inflammation ^65^, we hypothesise that the short-term shifts in inflammation and immune populations, which we found to be partially reversible, reflect a hormetic stress response that precedes long-term adaptation. Conversely, in buccal samples, we observed a reduction in immune cells, potentially reflective of lower immune infiltration. Comparison with a five-day fasting study in metabolic syndrome patients ^66^ revealed overlapping increases in monocytes, terminally differentiated CD8^+^ T cells, but diverging trends in B cells and naïve T cells, likely reflecting differences in population and study duration. In our cohort, IL-17A decreased selectively in stimulated T cells of postmenopausal women (**Extended Data Figure 11**), suggesting fasting may dampen inflammatory pathways linked to bone loss. These findings highlight the complex impact of fasting on immune status and inflammation, as well as its interplay with hormonal state.

The microbiome also showed tissue-specific responses. Oral microbiota underwent rapid restructuring, with reduction in Veillonella spp. and increases in *Porphyromonas pasteri* - taxa associated with biofilm dynamics and periodontal health ^45–48^ (**Figure 5**). Gut compositional changes were slower, but functionally consistent with enhanced SCFA production and vitamin biosynthetic potential in high compliant individuals. Diversity moved in opposing directions - decreased richness in saliva, increased Shannon diversity in stool, consistent with favourable patterns for oral and gut health, respectively. Of note, SR1 (Absconditabacteriales), a poorly characterised group typically present at low oral abundance in the mouth, reached over >10% in some participants, and Saccharimonadaceae emerged as a multi-omic hub, emphasising the need for functional exploration of understudied taxa in dietary interventions.

Metabolomic profiling revealed complementary dynamics. Saliva showed increases in SCFAs and decreases in essential amino acids, while urine profiles suggested improved glucose handling, altered one-carbon metabolism, and enhanced ketogenesis, with reductions in glucose, betaine, and 3-methylhistidine, and increased acetate (**Figure 4f**). These findings resonate with recent systemic profiling of exercise, where one-carbon metabolites such as betaine were identified as key mediators of metabolic adaptation, suggesting shared pathways across lifestyle interventions ^67^.

Both extrinsic (ketogenic supplementation) and intrinsic (menopausal status) modifiers shaped responses. MCT supplementation selectively altered cervical methylation, urine metabolites, and microbial taxa, sometimes tempering fasting-induced changes (**Figure 6**), contrary to our hypothesis that it may enhance the effects of fasting. Menopause strongly influenced baseline cervical methylation biomarkers and modulated immune and microbial responses, underscoring the importance of tailoring prevention strategies by hormonal state (**Extended Data Figure 11**).

Our integrative analyses connected these observations into systemic networks (**Figure 7**, **Extended Data Figures 14, 15**). Fat mass and ImmuneAge emerged as multi-omic hubs, connecting immune, microbial, and clinical features. Salivary Saccharimonadaceae correlated with buccal *LINE1* methylation, CRP, haematocrit, and epidermal thickness, suggesting an axis linking oral microbiota, inflammation, iron homeostasis, and epithelial integrity. Iron parameters correlated with VO_2_peak, BMI, creatinine, uric acid, and TNF-producing cytotoxic T cells, reinforcing the interplay of iron metabolism, inflammation, and systemic physiology. These cross-system associations exemplified how fasting engages coordinated networks rather than isolated pathways.

Our work extends beyond prior longitudinal omics cohorts (iPOP ^20,68^, the Pioneer 100 Wellness Project ^69^, a study by Marabita et al. 2022 ^19^) and intervention studies such as CALERIE™ ^70^ and METHOD ^71^, which have profiled longitudinal omics or examined lifestyle changes (e.g., caloric restriction), but often in few tissues. Here, repeated profiling across multiple non-invasive tissues, complemented by skin biopsies and clinical measurements, revealed tissue-specific responsiveness and systemic coordination of intervention effects, and notably a greater responsiveness of methylation biomarkers in epithelial samples. Together with our parallel smoking study, this dataset provides a benchmark for designing and interpreting health-promoting interventions and complements ongoing trials such as LIFE Tirol: Lasting health through IF, emotional fitness and exercise in Tirol, ISRCTN83248265).

## Limitations of the study

include the absence of a no-intervention control group, precluding definitive causal inference. We mitigated this by including compliance in our study design, performing a modified intention to treat analysis, as well as exploring within-person longitudinal changes. However, future studies should explore cross-over or run-in designs (e.g., LIFE, ISRCTN83248265). As the focus of this study was to quantify the health effect of a simple intervention (IF) rather than a specific diet, dietary intake was not systematically monitored to alleviate participant burden, though fasting adherence was validated against BMI and fat mass. Promising approaches to capture dietary intake through artificial intelligence tools (e.g., https://www.codiet.eu) may be considered in the future. Activity levels captured via wearables did not show major changes (**Extended Data Figure 1i**), but confounding through changes in physical activity cannot be excluded. Medication use was recorded at baseline but limited information was available longitudinally. While more detailed assessment and controlling for medication may be valuable in future studies, participants in the current study were predominantly healthy and only a small fraction (19/114, 16.67%) reported regular use of medication, predominantly antihypertensives, at baseline. The cohort was predominantly composed of Central European women, limiting generalisability. VO_2_peak has been described as a strong predictor of longevity ^72,73^, but it may be influenced by motivation ^74^, thus VO_2_peak increases with relation to IF should be confirmed in future studies. While our study features some dropouts, features were not predicted by baseline characteristics, suggesting lower risk of bias. Importantly, sensitivity analyses confirmed our results were robust to different modelling approaches (categorical, ordinal, or continuous time).

A key strength of this work is its breadth - deep, repeated, multi-tissue profiling in the same individuals, rigorous mixed-effects modelling with FDR correction, sensitivity analyses addressing dropout and modelling assumptions, and a public discovery app (https://eutops.github.io/lifestyle-atlas/docs/explore/app.html).

In conclusion, IF induced both systemic and tissue-specific changes, spanning iron metabolism, epithelial methylation, immune composition, metabolomic pathways, and microbiota changes. These changes converged on interconnected axes of iron homeostasis, adiposity, inflammation, and epithelial barrier integrity, offering insights into how fasting may modulate health trajectories. By integrating multiple omic layers in the same individuals, this study provides a benchmark resource for refining biomarkers of dietary and lifestyle interventions. More broadly, it highlights the potential of longitudinal, multi-tissue profiling to uncover coordinated physiological networks. As similar datasets accumulate, these approaches will inform biomarker development, reveal shared pathways across interventions such as diet and exercise, and may ultimately guide personalised strategies to extend human healthspan.

## Methods

### Study overview

This study encompasses the TirolGESUND intermittent fasting arm (NCT05678426). The TirolGESUND study has received ethical approval by the Ethics Committee of the Medical University of Innsbruck (Ethikkommission der Medizinischen Universität Innsbruck 1391/2020, 18.01.2021), Austria. The study is described in detail in a study and resource description paper ^17^. In brief, healthy participants aged 30-60 who had a BMI between 25 and 35 kg/m^2^ and did not suffer from hypo- or hyperthyroidism were recruited to participate in the study. All participants were healthy and free from current or former malignant, cardiometabolic, or psychiatric disease, and were not pregnant. Participants consented at baseline and completed several follow-up visits (month 2, 4, and 6, with optional month 12 and 18 follow-ups not included in the current analysis). Clinical measurements, samples, and questionnaire data were collected during visits or via digital tools at home.

### Study measures

Participants in the intermittent fasting followed a “time restricted eating” regime described by de Cabo and Mattson ^4^, guided by a dietician. A step-wise induction was instructed, whereby food intake was restricted to 10 h/day on 5 days a week in week 1, 8 h/day for 5 days in week 2, and subsequently 8 h on 7 days per week for the remainder of the study duration. Participants were also given the option to restrict food intake further to 6 h per week. Participants were randomised at baseline to receive a ketogenic supplement (Kanso MCTfiber, medium-chain triglyceride fiber sachet) or not to explore the beneficial effect of ketosis reinforcement, based on its suggested benefits in fasting ^75^, using a menopause- and BMI-stratified block randomisation strategy (blocks of 4) to ensure balanced groups. The ketogenic group received a powdered supplement (Kanso MCTfiber, medium-chain triglyceride fiber sachet) that could be added to food or diluted in water. Recommended use was the use of 1 sachet per day (10 g MCT) during weeks 1 and 2 and two sachets per day (20 g MCT) thereafter for the rest of the study period.

Each participant was assigned a personal coach who provided support and guidance and recorded compliance. Regular contact between the participant and coach, with a minimum recommendation of once a week, was encouraged. Where participants declined a coach, or no regular contact was maintained, compliant days per week since the last visit were recorded by study doctors during visits. Participants were offered the opportunity to participate in optional exercise guidance, tailored to fitness levels, as well as the option to engage in three supervised exercise sessions throughout the study, focusing on resistance, endurance, and flexibility training.

### Clinical measurements

Functional clinical measurements were collected using spirometry and ergometry at the Institute for Sports Medicine, Alpine Medicine and Health Tourism (ISAG), University Hospital/Tirol Kliniken in Innsbruck, Austria as outlined in our resource description paper ^17^. For the ergometry, all participants performed an incremental cycle test to exhaustion, starting at 50W with an increase of 25W for every 90s, which provided information on exercise capacity at relative stages to maximum capacity. VO_2_peak was calculated according to the following formula: VO_2_peak = Watt_peak_ * 12 + 3.2 * kg body weight. Body composition measurements were recorded using bioelectric impedance analysis, as described in our resource paper ^17^. Routine blood biochemistry was conducted using fasted samples while haemograms were performed fasted or unfasted. Vascular features and subcutaneous and visceral fat were measured using sonography and pulse wave velocity measurements at the Department for Neurology at the Medical University of Innsbruck, Austria.

### Wearable data

Data from Garmin vivosmart 4 were obtained via the Garmin Connect SDK Health API. Daily summary statistics were extracted. To compute activity levels, we computed average steps in two week intervals, at baseline (day 0-14) and two weeks before each person’s next visit date (M2, M4, M6).

### Methylation profiling

Methylation profiles were generated using the Illumina HumanMethylationEPIC version 1 in a high-throughput manner using liquid handling robots. Samples from the same individual were kept on the same beadchip, where possible, but positions of samples and visits were randomised to minimise batch effects. Data were preprocessed using a standardise pipeline (https://github.com/chiaraherzog/eutopsQC; previously reported e.g. in Herzog et al. ^76^) and thorough quality control was conducted to ensure no sample swaps occurred, including sample type analysis and SNP analysis of longitudinal samples. Some sample swaps could be identified and resolved based on SNP profiling and date of sample taking. Epigenetic biomarkers were computed using previously described coefficients or packages, and, where developed by other groups, validated using the Biolearn package ^77^ (see also our resource paper ^17^).

### Immune cell profiling via flow cytometry

Peripheral blood samples were stored at room temperature for less than 24 h before isolation of peripheral blood mononuclear cells. Samples were stained and immune cell populations were profiled in detail by flow cytometry, as previously described ^17^.

### Development of an ImmuneAge classifier

We developed an immune population-based classifier of chronological age using all baseline data (M0) from the smoking cessation and intermittent fasting study arms as training data, with all follow-up data as test sets. All immune populations (normalised values of white blood cell and T cell stainings) from baseline were used as input training data, with chronological age as the value to be predicted. To construct predictors, we explored ridge, lasso, or elastic net penalisation (α=1, 0, or 0.5, respectively) for linear regression (family=”gaussian”) in the glmnet R package (version 4.1.8). This approach followed a similar approach used to train predictors for epigenetic clocks, as previously described ^29^. Elastic net penalisation (alpha=0.5) exhibited the highest correlation with chronological age in baseline (training) data (R=0.63, **Figure 4b**) and was selected for the final model. Coefficients are shown in **Figure 4a**. ImmuneAge was then calculated using the coefficients extracted from linear models trained with elastic net regularisation and longitudinally evaluated in follow-up samples. Coefficients are provided in **Extended Data Table 13**.

### Saliva and urine metabolome profiling using nuclear magnetic resonance profiling, metabolite annotation, and metabolomics pathway enrichment analysis

Saliva and urine metabolome data were generated using nuclear magnetic resonance profiling (see resource paper ^17^). Briefly, frozen urine and saliva samples stored at −80°C were thawed at room temperature, and the supernatant after sedimenting solid particles at 14,000 rpm for 10 min was subjected to spectral analysis using an Avance II 600 MHz 349 spectrometer (Bruker Biospin, Rheinstetten, Germany), equipped with a Bruker 5 mm CryoProbe Prodigy TCI probe head with Z-gradient. Raw data underwent automatic Fourier transformation, phase correction, and baseline correction. Topspin 3.2 (Bruker) was employed for data processing, while Chenomx Profiler (Chenomx Inc., version 8.1) and the Human Metabolome Database HMDB (Version 5.0) ^78^ were utilised for the annotation of NMR peaks to compounds with manual checks. Spectral concentration was converted into comma-separated values (CSV) using the Microsoft Excel format for multivariate statistical analysis. Raw values were normalised using Total Sum Scaling to account for differences in concentration. Metabolites were annotated into classes using RefMet ^79^ (R package version 1.0.0). Metabolomics pathway enrichment analysis was conducted using the MetaboAnalystR package ^80^ (version 4.0.0). Significant metabolites for saliva metabolome (overall), urine metabolome (effect of time or modulation by compliance) were assessed using the SMPDB library, and enrichment scores (observed/expected) at p<0.05 were visualised.

### Analysis 16S rRNA gene amplicons

DNA isolation, dual barcode sequencing ^81^ of the V4 variable region of the 16S rRNA gene and inference of amplicon sequence variants (ASVs) with DADA2 ^82^ was done as described in our resource paper ^17^ for all stool and saliva samples from the smoking cessation and IF study arms collected within the TirolGESUND study. Automated taxonomic assignments done with the SINA classifier ^83^ (version 1.7.2) against SILVA database SSU Ref NR 99 (release 138.1) and Kraken2 ^84^ against the standard Kraken database (version 2.1.3) and manually curated as described in the parallel smoking cessation intervention study ^61^. Spurious ASVs that did not reach a relative abundance over 0.25 in any sample ^85^ were removed from the TirolGESUND saliva and stool microbiome datasets and only samples from the Intermittent Fasting study arm were retained for downstream analyses in R. Hellinger transformed ASV- and family-level count data (vegan version 2.6-4) was used for principal component analysis with FactoMineR ^86^ (version 2.9). Biplots and the significance of the association of the top three principal components with selected covariates were visualised with factoextra ^87^ (version 1.0.7) and ComplexHeatmap ^88^ (version 2.12.1), respectively. The count data was rarefied to the minimum sequence depths among the stool or saliva samples and the alpha diversity indices Chao1 and Shannon were calculated with ampvis2 ^89^ (version 2.7.36) wrapper for vegan. Changes over time in alpha diversity in the high compliance group was evaluated comparing the median indices at months 2, 4, 6 with those at baseline (paired two-sided Wilcoxon signed-rank tests, ggpubr version 0.6.0). Functional predictions were done for all ASVs and associated (non-rarefied) count data with PICRUST2 ^90^ (version 2.5.2). Microbiome features used for longitudinal linear mixed effect modelling as described were first transformed. Centred log-ratio (CLR) transformation was applied on the ASV-level count data after adding a small pseudo-count using the microbiome R package (version 1.26.0). Family-level and predicted KEGG enzyme count data were normalised by Total Sum Scaling (TSS).

### Statistical analysis

#### Power calculation

The study was powered to detect an initial estimate of a clinically relevant effect in DNAme indices (e.g., WID-BC), assuming a standard deviation of 0.44 (based on data from the FORECEE validation set), a statistical power of 94% and a significance level of 5%. As there was no preliminary data on the longitudinal dynamics of DNAme indices (e.g., WID scores) or other molecular dynamics quantified during the study, the TirolGESUND study also serves as a pilot study to assess effect sizes for future studies.

#### Data analysis approach

We conducted both a modified intention to treat (mITT) and per protocol (PP) analysis, leveraging compliance categories (**Figure 1**) on complete cases. We motivate the use of complete cases and mITT by our primary intention to study the impact of intermittent fasting on longitudinal biomarkers within a given individual. This approach allowed us to use all available longitudinal data from participants.

To do so, we performed both paired two-sided Wilcoxon tests as well as linear mixed-effects models that allowed us to additionally adjust for covariates (e.g., age or BMI at baseline) and explore the interaction between time and compliance group. For ease of use for multi-omics data integration, we leveraged the MultiAssayExperiment format ^91^ (MultiAssayExperiment R package version 1.35.3).

#### Modified intention to treat and per protocol analysis using Wilcoxon tests

Absolute clinical changes were evaluated using paired two-sided Wilcoxon tests compared to baseline on complete cases for each feature (**Extended Data Table 2**), while differences in the change from baseline by compliance group or intervention group (high compliance individuals only) were compared using unpaired two-sided Wilcoxon tests (**Extended Data Tables 3, 4, respectively**). Longitudinal changes compared to baseline in the high compliance group only were computed using paired two-sided Wilcoxon tests compared to baseline (**Extended Data Table 5**).

#### Modified intention to treat and per protocol analysis using linear mixed-effects models

In a second approach to assess the impact of time, compliance, or intervention group on clinical and omic features, we fitted linear mixed effects (LME) models to fully leverage the longitudinal data, adjust for age at consent, and estimate the interaction between time since baseline and compliance. Briefly, each feature was normalised (thereafter referred to as ‘value’) and three models were run on the data. All models used fixed slopes and random intercepts and visit was coded as a categorical factor to allow for diverse temporal dynamics, and models were computed using the R package lme4 (version 1.1.35.1). Model a (“Basic model”) explored the impact of time on the variable overall, adjusting for age and BMI at consent impact of time. The visitId estimates were extracted from the following model: value ~ age at consent + bmi at consent + visitId + (1|subjectId). Model b (“Interaction model”) explored the interaction between time and high compliance, accounting for age and BMI at baseline, and extracted estimates from the time and compliance interaction term from the following model: value ~ age at consent + bmi at consent + visitId*compliance + (1|subjectId). Finally, model c (“Ketogenic supplement model”) explored the impact of time and intervention allocation in highly compliant individuals. Values of highly compliant individuals were adjusted for age and BMI at consent and the interaction term for time and intervention group was extraced from the following model: value ~ age at consent + bmi at consent + visitId*interventionId + (1|subjectId).

All models used the maximum likelihood (ML) estimation method. p values obtained from Satterthwaite’s degrees of freedom approximation for models run were extracted using the lmerTest R package (version 3.1.3). Zero inflation in the context of microbiome data was handled by removing spurious ASVs and CLR transformation of the ASV-level count data.

#### Sensitivity analysis of linear mixed-effects model time coding

Time was by default coded as a categorical (factor) variable (M0, M2, M4, M6, with M0 as the baseline value) to allow for non-gradual, non-linear, and non-unidirectional effects, as well as allow for simple visualisation of effects at each visit. To perform a sensitivity analysis on the impact of coding time either as an ordinal (M0<M2<M4<M6) or continuous variable (0, 2, 4, 6), we ran models with these alternative codings of time and compared obtained estimates and p values for time or the interaction of time with high compliance.

#### Variance decomposition

To estimate between- and within-individual variation, we fitted the following linear mixed-effects model using the R package lme4 (version 1.1.35.1): value ~ visitId*compliance + age_at_consent + bmi_at_consent + interventionId + (1 | subjectId), where value represented the normalised feature. The intraclass correlation coefficient (ICC) was computed as the proportion of total variance explained by the subject structure, i.e. V_random_/V_total_, as extracted from linear mixed effects models using the get_variance function from the insight R package (version 0.19.8). For 31 out of the 49 saliva metabolites no ICC could be calculated due to singular fits of the random effect term (1|subjectId). Singular fits indicate that the model could not reliably estimate variance attributable to the subject structure, suggesting that the random effect variance is close to zero or the model is overparameterized for the majority of the saliva metabolome features. The urine metabolome exhibited a slightly higher ICC, indicating it may be in part regulated by metabolic processes, although the overall ICC was still low compared to other omic measurements.

#### Association of features with menopause and interaction of menopause and compliance

The association of menopause with features was explored using linear mixed-effects models with fixed slopes and random intercepts. Model a) extracted the p value for menopause from a model accounting for age and BMI at baseline, visit, and intervention group (value ~ age at consent + bmi at consent + visitId + menopause + (1|subjectId)). Model b) extracted the p value for the interaction of menopause and visit in a model accounting for age and BMI at baseline and intervention group (value ~ age at consent + bmi at consent + visitId*menopause + (1|subjectId)). Models were run using lme4 (version 1.1.35.1) in R, and p values were extracted using the lmerTest R package (version 3.1.3).

#### Association of baseline characteristics with change in BMI

The association of baseline characteristics and change in BMI at month 6 compared to baseline was computed for each feature separately using Spearman correlation for continuous or Kruskall-Wallis test for categorical values. P values were adjusted using Holm-Bonferroni correction.

#### MEFISTO

We applied MEFISTO ^57^ for multi-omic, time-aware factor analysis on our data. A probabilistic model was trained on per ome normalised and z-scaled, centred data grouped by intervention from the blood haemogram, T cell and white blood cell staining, metabolomes, microbiomes (families) and epigenetic biomarkers using the R MOFA2 package (version 1.13.0), relying on python’s mofa2py (parameters: maximum 11 dimensions, 200 training iterations, start_opt=50, opt_freq=50). The association across the two study groups of the resulting 15 learned latent factors with different covariates was further examined with correlation analyses (Pearson correlation coefficient or Kruskal-Wallis test), the association with the intervention in the different compliance groups with paired two-sided Wilcoxon tests.

#### Repeated measures correlation analysis

We explore longitudinal associations of omic features using the rmcorr_mat function from the rmcorr R package (version 0.6.0) on a matrix in which rows represented observations and columns represented normalised features. We extracted cross-omic features (i.e., retaining only significant correlations at FDR<0.05 (within each feature)) that spanned across different assays or sample types.

#### Adjustment for multiple testing

We accounted for multiple testing via false discovery rate (FDR) correction. FDR correction was conducted within each data layer (e.g., clinical, biomarkers, microbial families) or individual measure (repeated measures correlation). P values that remained below<0.05 after FDR correction within ome (Wilcoxon tests, paired plots, heatmaps - bold significance stars and labels in heatmaps) or across all comparisons (repeated measures correlation analysis) were considered statistically significant. Additionally, for the effect of time, interaction of time and compliance, and interaction of time and allocated intervention, we also employed permutation testing as a complementary approach to explore our findings. One hundred (100, n) iterations of models were run, randomly permuting visitId (basic and interaction models) or intervention group (ketogenic supplement models), and p values were extracted. Expected p values from permutation analyses (p_expected_) were then compared to observed p values (p_observed_). If the observed p value was significantly lower than the expected p value at p<0.05, derived from the following equation:

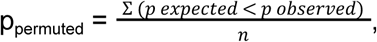

the result was considered significant (grey significance stars and italic labels in heatmaps).

### Data visualisation

Data were visualised in R. Boxplots, scatter plots, and bubble plots, or where otherwise specified below, were visualised using ggplot2 (version 3.5.0). Principal component analysis was conducted using the FactoMineR ^86^ T package (version 2.9) and coordinates extracted for screeplots, while biplots were visualised using factoextra ^87^ (version 1.0.7). Heatmaps were generated using ComplexHeatmap ^88^ (version 2.12.1). Tree diagrams for immune populations were generated by transforming hierarchical data into a tree format using the tidytree package (version 0.4.6) and visualising the tree using the ggtree package (version 3.10.1). The Chord diagram for cross-ome correlations were visualised using the circlize R package (version 0.4.16). Cross-ome network diagrams were generated using the igraph R package (version 2.0.2), filtering first connections of selected biomarkers. Long format data were transformed into a network using the graph_from_edgelist function. Node colours were set by assay type while edge colour and size was plotted by correlation sign and strength.

## Acknowledgements

We thank all participants of the TirolGESUND study and individuals involved in the administration of the study. We thank our research nurses Gabi Hilber, Barbara Rovara and Birgit Kröss. We thank Maria Meister and her lab team for the assistance with sample processing. We thank the nutrition and dietetics team, Ute Pichler, Markus Augschöll, David Ebner and Heidi Spildenner who performed dietetic counselling and bioelectric impedance analysis. We thank the the vascular measurement team Silvia Komarek, Benjamin Dejakum, (all Department of Neurology, Medical University of Innsbruck; VASCage - Research Centre on Vascular Ageing and Stroke, Innsbruck), Alex Messner (Department of Pediatrics II, Medical University of Innsbruck), Bernhard Winder (Department of Vascular Surgery, Feldkirch Hospital, Feldkirch, Austria), and Johannes Nairz (VASCage-Research Centre on Vascular Ageing and Stroke, Innsbruck; Department of Pediatrics II and III, Medical University of Innsbruck). We thank Dr. Schär AG / SPA (Winkelau 9, 39014 Burgstall (BZ) Italy) for providing the Kanso MCTfiber. We thank the psychology students Minou Mohraz, Olaya Roces Sanchez, Rosa Huber, Carina Zeitler, Anne Harbring, Rosa Ottmann, Caroline Siebert, Valeria Kovalchuk, Elvira Galeazzo, Lina Förster, Pauline Raßbach, Sonja Breu, Anna Zellmer, Elif Algül, Melisa Amin, Johanna Haessler, Lilli Schulz, Valerie Nickel, Lea Rosenstock for coaching and supporting the volunteers. Chiara Herzog’s primary affiliation is now King’s College London.

## Author contributions [CRediT statement]

**Chiara MS Herzog:** conceptualisation, methodology, formal analysis, data curation, writing - original draft, visualisation, supervision, project administration

**Charlotte Dafni Vavourakis**: methodology, formal analysis, visualisation, data curation, writing - original draft

**Bente Theeuwes:** formal analysis, visualisation, data curation, software, writing - review and editing

**Elisa Redl:** investigation, project administration, writing - review and editing

**Christina Watschinger**: investigation, project administration, writing - review and editing.

**Gabriel Knoll**: investigation, data curation, writing - review and editing

**Magdalena Hagen**: investigation, data curation, writing - review and editing

**Andreas Haider**: investigation, validation, writing - review and editing

**Hans-Peter Platzer**: investigation, data curation, writing - review and editing

**Umesh Kumar**: investigation, data curation, writing - review and editing

**Sophia Zollner-Kiechl**: investigation, data curation, writing - review and editing

**Maria Cavinato**: investigation, data curation, writing - review and editing

**Pidder Janssen-Duerr**: supervision, writing - review and editing

**Matthias Schmuth**: investigation, data curation, writing - review and editing

**Maximilian Lammer**: investigation, writing - review and editing

**Michael Knoflach**: investigation, data curation, writing - review and editing

**Verena Lindner**: investigation, supervision

**Anna Wimmer**: investigation, supervision

**Tobias Greitemeyer**: investigation, supervision

**Peter Widschwendter**: investigation, supervision

**Sonja Sturm**: investigation, supervision, data curation, writing - review and editing

**Hermann Stuppner**: resources, writing - review and editing

**Birgit Weinberger**: supervision, resources, writing - review and editing

**Alexander Moschen**: resources, project administration, writing - review and editing

**Alexander Höller**: supervision, resources

**Wolfgang Schobersberger**: supervision, resources

**Martin Widschwendter**: conceptualisation, project administration, resources, writing - original draft, supervision, funding acquisition

## Data availability

Data availability is described in detail in our resource paper ^17^ and links to digital object identifiers of datasets will be provided on the Tyrol Lifestyle Atlas Data Portal (https://eutops.github.io/lifestyle-atlas/). In brief, methylation and microbiome data are deposited on EGA under accession codes EGAS00001007840 and EGAS00001007842, respectively. Other assay data are deposited on Zenodo in tabular format. Anonymised data will be available openly after a one year embargo period. Study data can be explored using the Tyrol Lifestyle Atlas Data App (https://eutops.github.io/lifestyle-atlas/explore/).

## Code availability

Code for analyses is provided on github under (https://github.com/chiaraherzog/MultiOmics_IntermittentFasting).

