## Extended Data for "Systemic multi-omic remodelling underlies health benefits of intermittent fasting"

**Extended Data Tables** 4

Extended Data Table 1. Participant characteristics of participant characteristics at baseline by allocated study arm. 4

Extended Data Table 2. Modified intention to treat analysis comparing baseline values to month 2, 4, and 6 (paired two-sided Wilcoxon test) in individuals who completed the study. 4

Extended Data Table 3. Comparison of changes from baseline in the higher compliance versus lower compliance group (unpaired two-sided Wilcoxon test at month 2, 4, and 6) in individuals who completed the study. 4

Extended Data Table 4. Comparison of changes from baseline in the I versus K study arm, within highly compliant individuals (unpaired two-sided Wilcoxon test at month 2, 4, and 6). 4

Extended Data Table 5. Per protocol analysis comparing baseline values to month 2, 4, and 6 (paired two-sided Wilcoxon test) in highly compliant individuals who completed the study. 4

Extended Data Table 6. Modified intention to treat analysis using linear mixed effects models to compare baseline values to month 2, 4, and 6, adjusting for age at baseline and BMI at consent in individuals who completed the study. 5

Extended Data Table 7. Linear mixed effects models exploring the interaction of time and high compliance in individuals who completed the study, adjusting for age and BMI at consent. 5

Extended Data Table 8. Linear mixed effects models exploring the interaction between time and study group (I versus K) in highly compliant individuals who completed the study (month 2, 4, and 6). 5

Extended Data Table 9. Impact of menopause on features overall, derived from linear mixed-effects modelling. 5

Extended Data Table 10. Modulation of effects by menopausal status in linear mixed-effects modelling. 5

Extended Data Table 11. Association of baseline features with change in BMI over the intervention. 5

Extended Data Table 12. Significant features for repeated measures correlation. Only features with FDR corrected p values < 0.05 are considered. 5

**Extended Data Figures** 6

Extended Data Figure 1. Details on clinically reported effects, MCT supplement intake, and health-related quality of life questionnaire results. 6

Extended Data Figure 2. Principal component analysis of most personally variable features. 8

Extended Data Figure 3. Clinical changes by compliance group. 9

Extended Data Figure 4. Changes in absolute values of iron-related blood features. 10

Extended Data Figure 5. Principal component analysis details and longitudinal changes in selected composite epigenetic biomarkers. 12

Extended Data Figure 6. Immune cell variability and changes over time in different populations. 13

Extended Data Figure 7. T cell cytokine production is altered over the course of intermittent fasting. 15

Extended Data Figure 8. Principal component analysis on saliva and urine metabolome data. 17

Extended Data Figure 9. PCA on microbial ASVs and ASVs associated with age and menopausal status. 18

Extended Data Figure 10. Alpha-diversity indices and family-level taxonomic composition of the microbiomes. 19

Extended Data Figure 11. Features associated with menopause and treatment interaction with menopause. 20

Extended Data Figure 12. Defective NK cells at baseline predict ΔBMI at month 6. 21

Extended Data Figure 13. Sensitivity analysis of models with regards to coding the time variable as a categorical (factor), ordinal, or numeric for basic models (a, c-f) or interaction

models (b, g-j). 22
Extended Data Figure 14. Integrative omic MEFISTO analyses. 24
Extended Data Figure 15. Repeated measures correlation results and examples for ageing--related biomarkers and most connected features. 25
Extended Data Figure 16. Repeated measures correlation network of iron-related metabolism. 27
**Supplementary Data 28**
Supplementary Figure 1. Data for fat mass illustrating impact of compliance and intervention, as summarised in Figure 2k. 28

#### Extended Data Tables

**Extended Data Table 1. Participant characteristics of participant characteristics at baseline by allocated study arm.**
MCT denotes multi-chain triglycerides.

| Characteristic | Intermittent fasting n = 45 | Intermittent fasting plus MCT n = 51 |
|---|---|---|
| **Age at consent**, Mean (SD) | 48.5 (6.3) | 45.8 (8.6) |
| **BMI at consent** (kg/m²), Mean (SD) | 29.3 (2.9) | 29.8 (2.9) |
| **Postmenopausal**, n (%) | 15 (33) | 17 (33) |
| **Dietary preference**, n (%) | | |
| normal | 38 (84) | 47 (92) |
| vegetarian/vegan/pescetarian | 1 (2.2) | 2 (3.9) |
| unknown | 6 (13) | 2 (3.9) |
| **Smoking history**, n (%) | | |
| Never smoker | 20 (44) | 33 (65) |
| Former smoker | 20 (44) | 16 (31) |
| unknown | 5 (11) | 2 (3.9) |
| **Weekly intense activity (min)**, n (%) | | |
| no intense activity | 23 (51) | 29 (57) |
| 30-149 min | 15 (33) | 14 (27) |
| ≥150 min | 2 (4.4) | 6 (12) |
| unknown | 5 (11) | 2 (3.9) |
| **Systolic blood pressure** (mmHg), Mean (SD) | 117.6 (14.4) | 115.5 (13.2) |
| **Diastolic blood pressure** (mmHg), Mean (SD) | 76.0 (8.6) | 76.5 (8.3) |
| **Total cholesterol** (mg/dL), Mean (SD) | 207.1 (27.9) | 200.5 (33.8) |
| **Triglycerides** (mg/dL), Median (IQR) | 90.0 (49.0) | 86.0 (47.5) |
| **HbA1c** (%), Mean (SD) | 5.4 (0.3) | 5.3 (0.3) |
| **Fasting glucose** (mg/dL), Mean (SD) | 81.7 (13.8) | 80.6 (14.4) |
| **Haemoglobin** (g/dL), Mean (SD) | 13.3 (0.8) | 13.1 (0.9) |

**Extended Data Table 2. Modified intention to treat analysis comparing baseline values to month 2, 4, and 6 (paired two-sided Wilcoxon test) in individuals who completed the study.**
Provided as .xlsx file.

**Extended Data Table 3. Comparison of changes from baseline in the higher compliance versus lower compliance group (unpaired two-sided Wilcoxon test at month 2, 4, and 6) in individuals who completed the study.**
Provided as .xlsx file.

**Extended Data Table 4. Comparison of changes from baseline in the I versus K study arm, within highly compliant individuals (unpaired two-sided Wilcoxon test at month 2, 4, and 6).**
Provided as .xlsx file.

**Extended Data Table 5. Per protocol analysis comparing baseline values to month 2, 4, and 6 (paired two-sided Wilcoxon test) in highly compliant individuals who completed the study.**
Provided as .xlsx file.

**Extended Data Table 6. Modified intention to treat analysis using linear mixed effects models to compare baseline values to month 2, 4, and 6, adjusting for age at baseline and BMI at consent in individuals who completed the study.**
Each variable was normalised to provide standardised estimate sizes. Time was coded as a categorical variable (visitId). Provided as .xlsx file.

**Extended Data Table 7. Linear mixed effects models exploring the interaction of time and high compliance in individuals who completed the study, adjusting for age and BMI at consent.**
Each variable was normalised to provide standardised estimate sizes. Time was coded as a categorical variable (visitId). Provided as .xlsx file.

**Extended Data Table 8. Linear mixed effects models exploring the interaction between time and study group (I versus K) in highly compliant individuals who completed the study (month 2, 4, and 6).**
Each variable was normalised to provide standardised estimate sizes. Time was coded as a categorical variable (visitId). Provided as .xlsx file.

**Extended Data Table 9. Impact of menopause on features overall, derived from linear mixed-effects modelling.**
Each variable was normalised to provide standardised estimate sizes. Time was coded as a categorical variable (visitId). Provided as .xlsx file.

**Extended Data Table 10. Modulation of effects by menopausal status in linear mixed-effects modelling.**
Each variable was normalised to provide standardised estimate sizes. Time was coded as a categorical variable (visitId). Provided as .xlsx file.

**Extended Data Table 11. Association of baseline features with change in BMI over the intervention.**
Provided as .xlsx file.

**Extended Data Table 12. Significant features for repeated measures correlation.** Only features with FDR corrected p values $< 0.05$ are considered.
Provided as .xlsx file.

#### Extended Data Figures

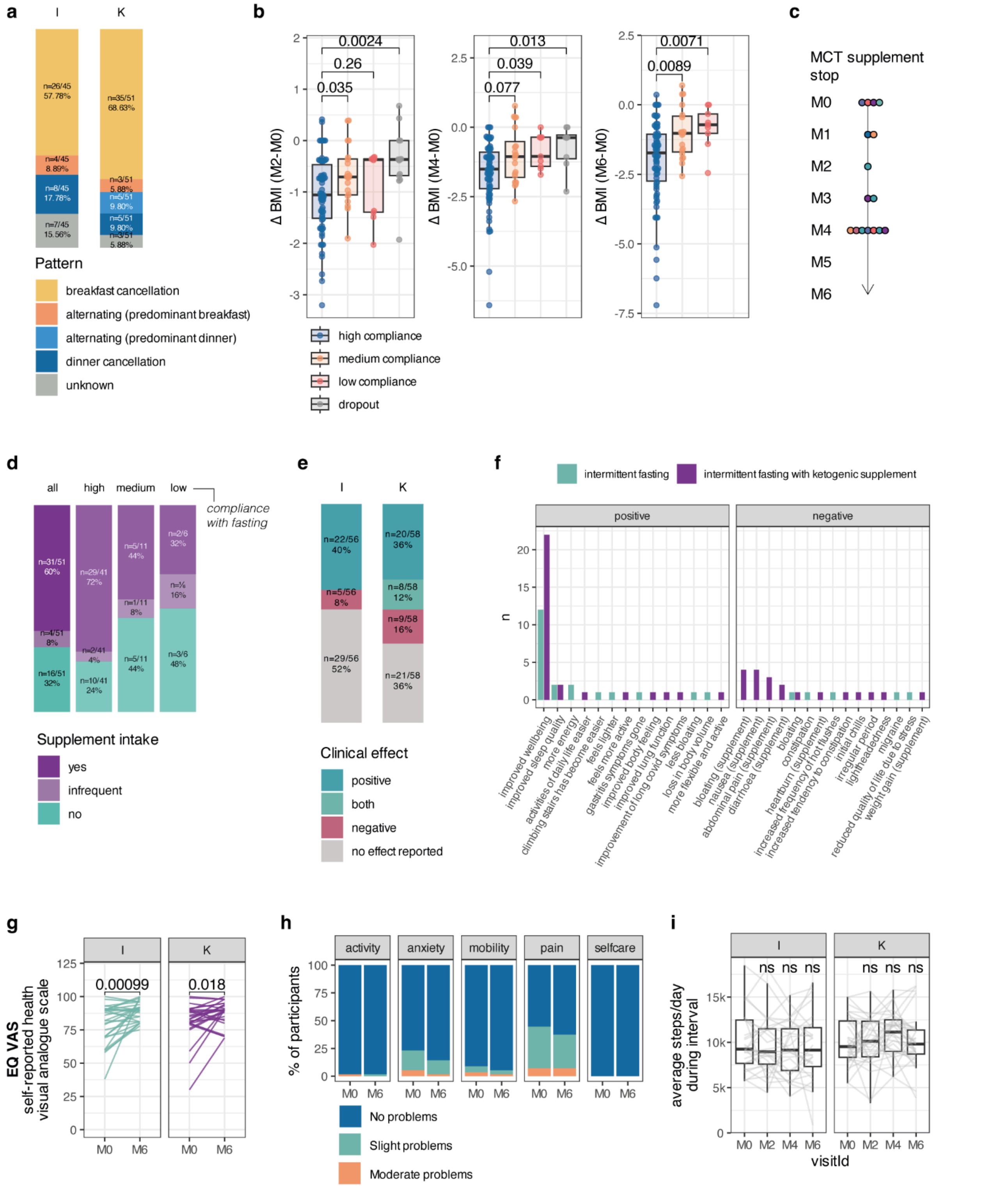


**Extended Data Figure 1. Details on clinically reported effects, MCT supplement intake, and health-related quality of life questionnaire results.**

**a** Fasting pattern in the two study arms in individuals who completed the study. **b** Change in BMI from baseline (Δ BMI) by compliance and intervention group. **c** Time at which supplement intake was stopped in the 16 individuals who reported not to use the MCT supplement. Each dot represents one individual. **d** Compliance with the MCT supplement intake in the intermittent fasting plus MCT ketogenic supplement (K) study arm, restricting to individuals who completed the study. **e** Clinical effects reported by participants. **f** Details on reported clinical effects. (supplement) in brackets indicates specific symptoms reported to be associated with the supplement use. **g** EQ-VAS (visual analogue scale) comparing baseline and M6 in the two randomised study arms. p value is derived from a paired two-sided Wilcoxon test. **h** EQ-5D-5L scores at baseline and M6. **i** Average steps per day per participant from wearable device; intervals include baseline (first two weeks, M0) and thereafter two weeks prior to each visit (M2, M4, M6).

**Abbreviations:** I, Intermittent fasting group. K, Intermittent fasting plus multi-chain triglyceride (ketogenic supplement). MCT, multi-chain triglyceride. M0, M2, M4, M6: month 0, 2, 4, 6. EQ-VAS: EuroQoL (quality of life)-Visual Analogue Scale.

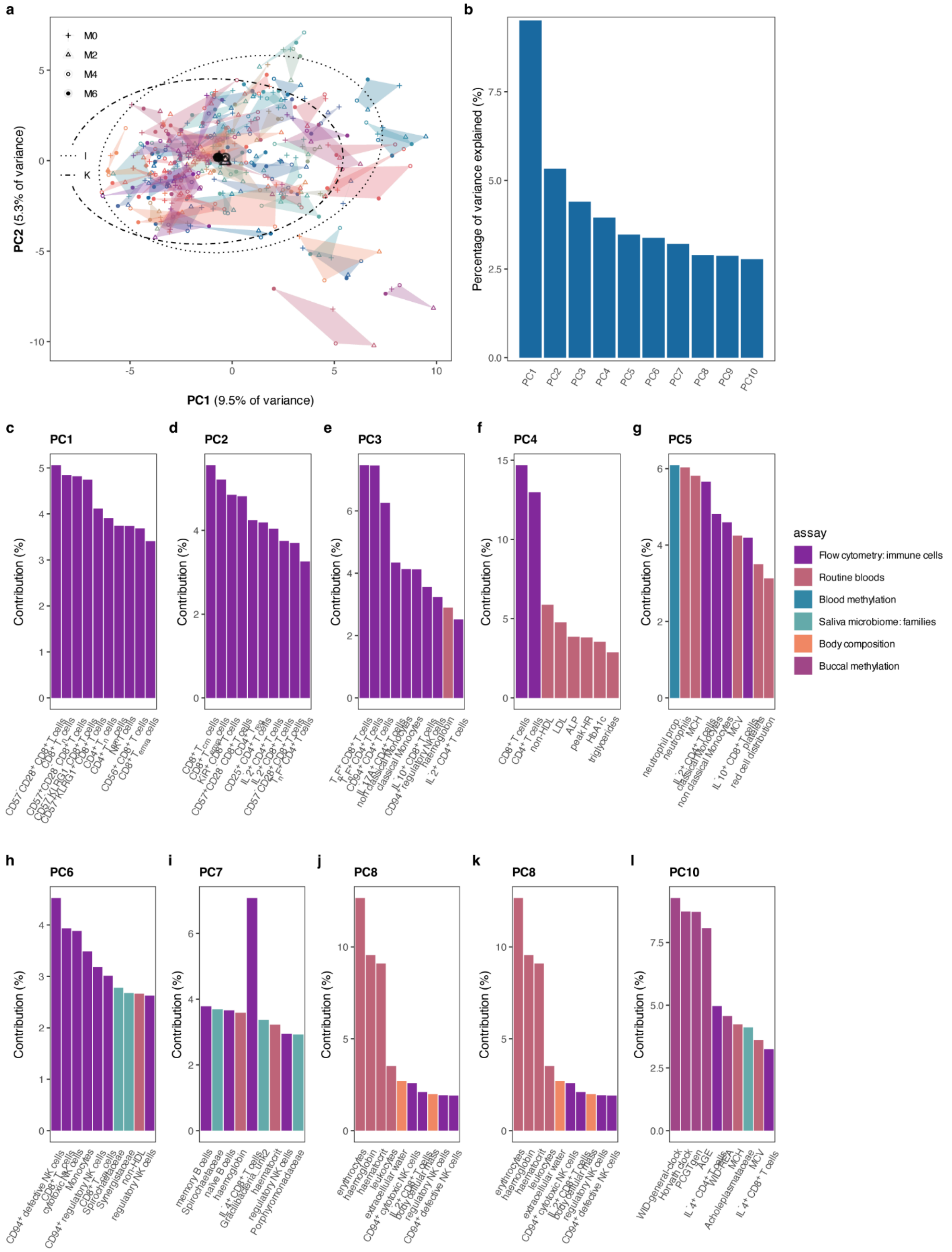


**Extended Data Figure 2. Principal component analysis of most personally variable features.**
**a** Individual trajectories for principal components 1 and 2. Shapes represent visits with large shapes indicating centroids for visits at month 0, 2, 4, and 6 in all individuals. Each hull represents one individual. Ellipses represent each intervention. **b** Scree plot of percentage of variance explained by each principal component. **c-l** Top 10 feature contributors for the first ten principal components, coloured by assay.

**Abbreviations:** M0, M2, M4, M6: month 0, 2, 4, 6. PC, principal component.

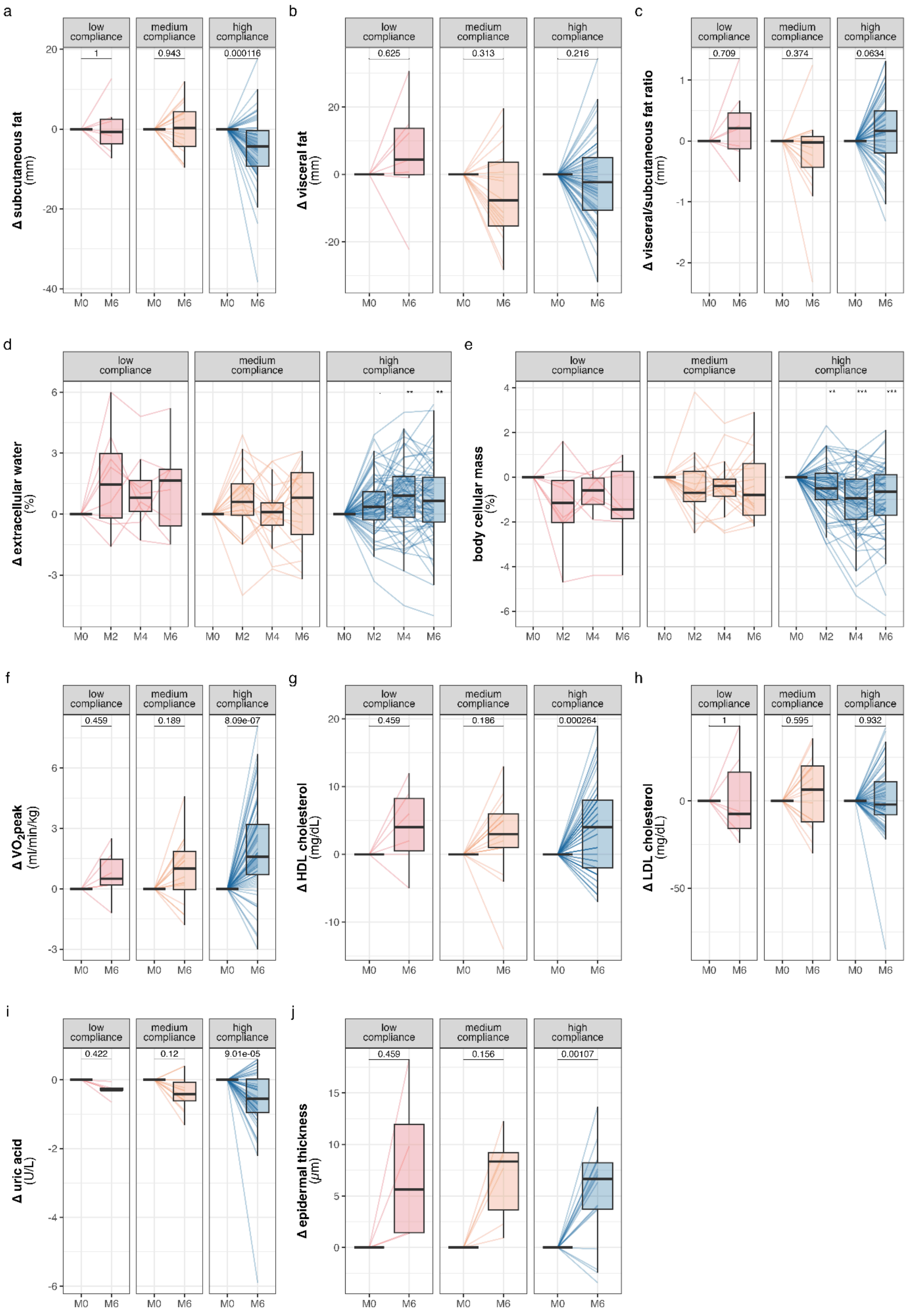


**Extended Data Figure 3. Clinical changes by compliance group.**
Change from baseline in **a** subcutaneous fat, **b** visceral fat, **c** visceral to subcutaneous fat ratio, **d** body cellular mass, **e** $VO_2$peak, **f** HDL cholesterol, **g** LDL cholesterol, **h** uric acid, and **i** epidermal thickness by compliance group. p values for paired comparisons are derived from paired two-sided Wilcoxon test, compared to baseline values, and all p values are FDR corrected within “ome” (all clinical and routine laboratory measurements) and timepoint.
**Abbreviations:** HDL, high-density lipoprotein. LDL, low-density lipoprotein.
Boxplots indicate median and interquartile range while whiskers indicate values +/- 1.5 interquartile range. Individual data points are shown where possible.

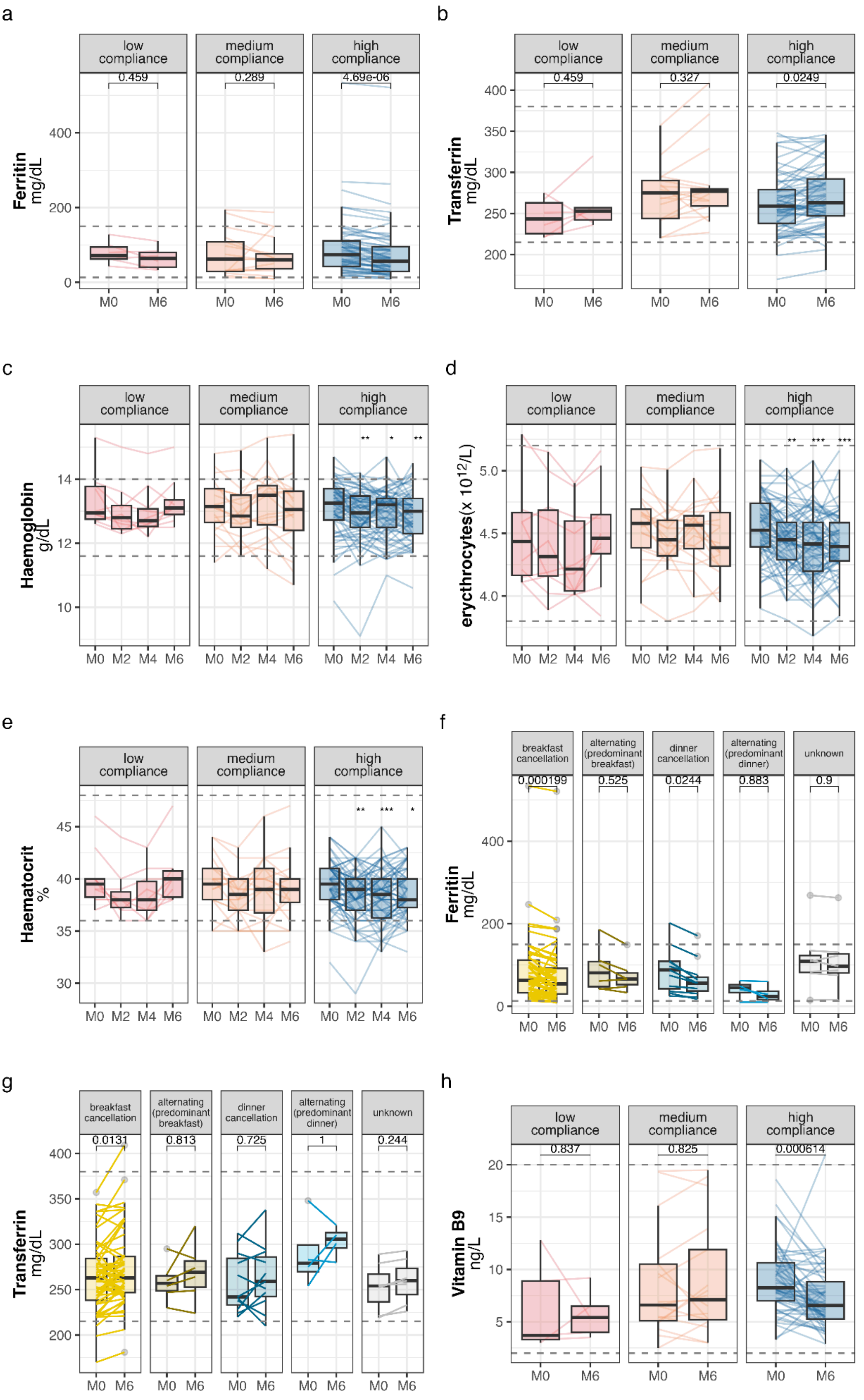


**Extended Data Figure 4. Changes in absolute values of iron-related blood features.**
**a** Raw values of ferritin, **b** transferrin, **c** haemoglobin, **d** erythrocytes, **e** haematocrit, and **h** vitamin B9 by compliance groups. **f** and **g** Changes to ferritin and transferrin by fasting pattern and intervention. Respective reference values are indicated with dashed lines. p values are derived

from paired two-sided Wilcoxon tests compared to baseline and FDR corrected within “ome” (all clinical and routine laboratory measurements) and timepoints.
Boxplots indicate median and interquartile range while whiskers indicate values +/- 1.5 interquartile range. Individual data points are shown where possible.

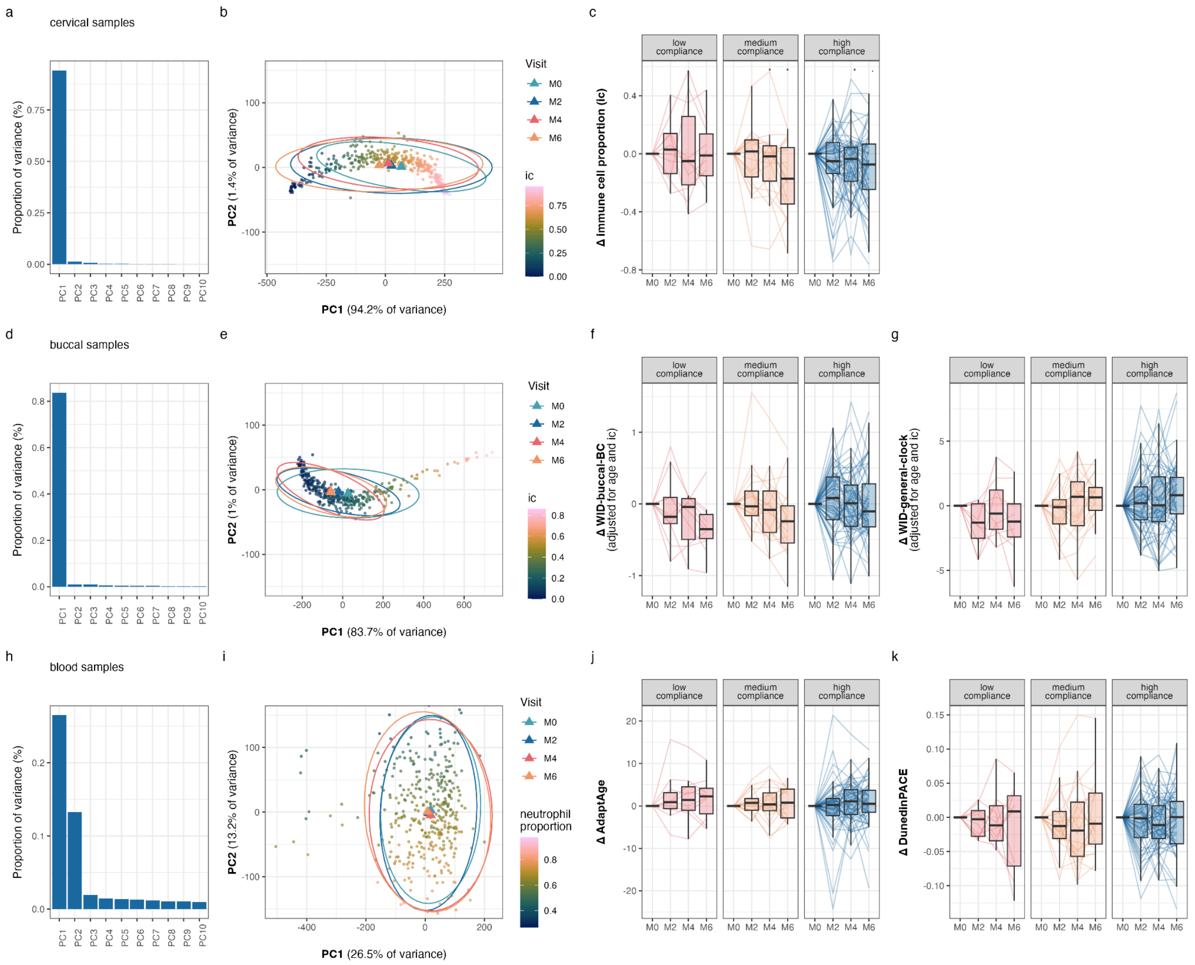


**Extended Data Figure 5. Principal component analysis details and longitudinal changes in selected composite epigenetic biomarkers.**

**a** Scree plot for top 5% variable CpGs in cervical DNA methylation data. **b** PC1 and PC2 for top 5% variable CpGs in cervical samples indicate a strong dependence on immune cell proportion and a gradual shift towards lower immune cell proportion over time (M0 to M6). **c** Change in immune cell proportion from baseline by compliance group. p values derived from paired two-sided Wilcoxon tests compared to baseline. **d** Scree plot for top 5% variable CpGs in buccal DNA methylation data. **e** PC1 and PC2 for top 5% variable CpGs in buccal samples indicate a strong dependence on immune cell proportion and a gradual shift towards lower immune cell proportion over time (M0 to M6). **f** Change in WID-buccal-BC or **g** WID-general-clock index in buccal samples from baseline by compliance group. p values derived from paired two-sided Wilcoxon tests compared to baseline. **h** Scree plot for top 5% variable CpGs in blood DNA methylation data. **i** PC1 and PC2 for top 5% variable CpGs indicate a limited dependence on time. **j** Change in AdaptAge and **k** DunedinPACE from baseline by compliance group. p values derived from paired two-sided Wilcoxon tests compared to baseline.

Boxplots indicate median and interquartile range while whiskers indicate values +/- 1.5 interquartile range. Individual data points are shown where possible. All p values were FDR corrected. If no p value is shown, p>0.05.

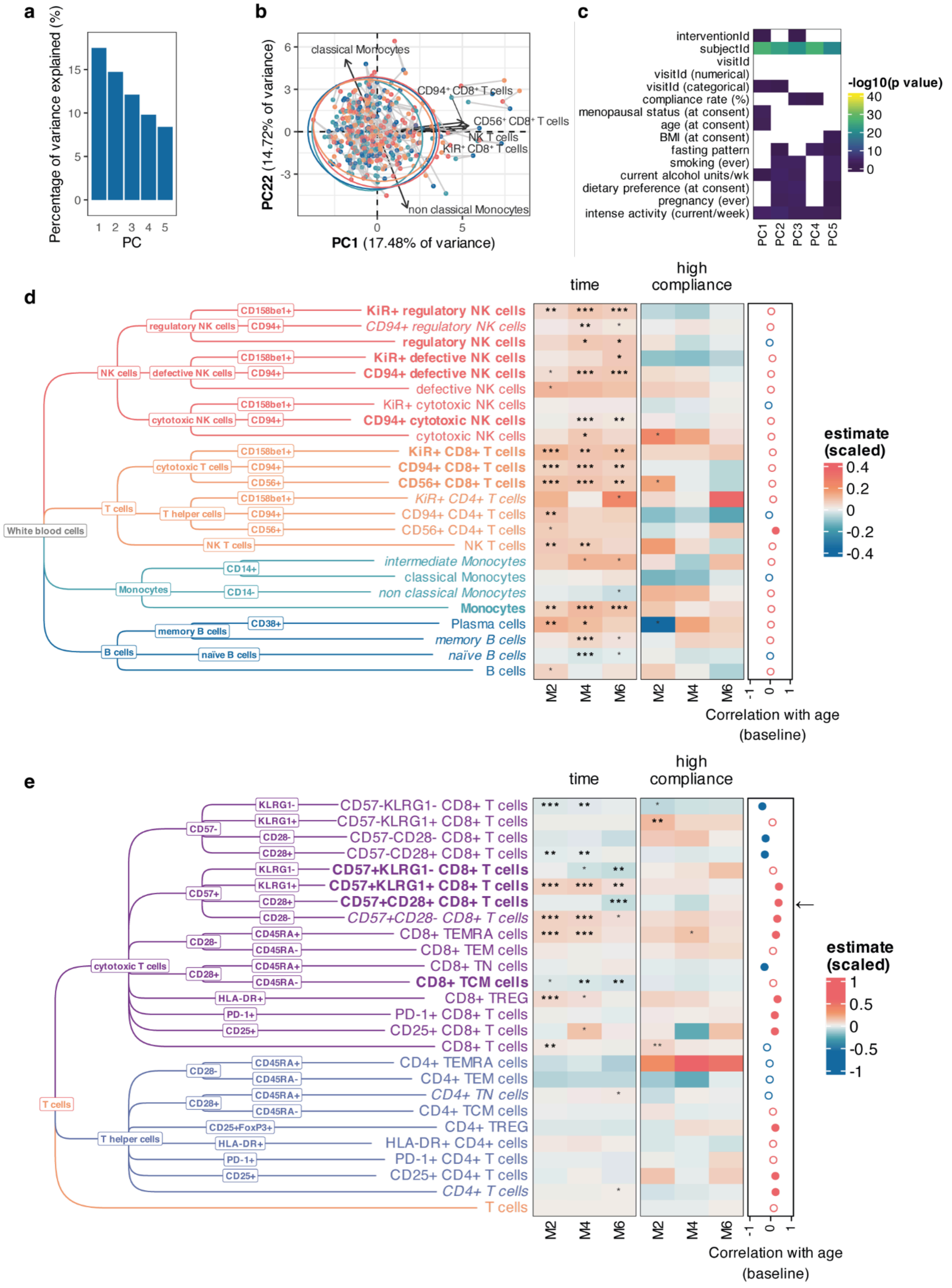


**Extended Data Figure 6. Immune cell variability and changes over time in different populations.**

**a** Scree plot showing the percentage of variance explained by the first five principal components in immune cells. **b** The first two principal components grouped by participant and visit. The top 5 contributors to the principal components are indicated. **c** Heatmap of p values for associations of principal components with participant and study characteristics. **d** Tree diagram for immune cell populations analysed using white blood cell staining and estimates from linear-mixed effects models impact of time (value ~ age at consent + bmi at consent + visitId + (1|subjectId)) and interaction between time and high compliance (value ~ age at consent + bmi at consent + visitId*compliance + (1|subjectId)). **e** Tree diagram for immune cell populations analysed with T cell staining and estimates from linear-mixed effects models impact of time (value ~ age at consent + bmi at consent + visitId + (1|subjectId)) and interaction between time and high compliance (value ~ age at consent + bmi at consent + visitId*compliance + (1|subjectId)), and time in immune cells. For d and e, *, **, *** indicate $p < 0.05$, $< 0.01$, and $< 0.001$, respectively. Values that remain significant after FDR correction are indicated in bold; row labels of significant features after FDRare shown in bold. Values that remain significant after permutation testing correction are indicated in grey; row labels of significant features at month 6 after permutation testing are shown in italic. Correlation with age at baseline values is indicated on the right hand side (positive or negative in red and blue, respectively). Significant correlations at $p<0.05$ are indicated with filled circles. Populations that go into the opposite direction as ageing over time in the study are indicated with arrows.

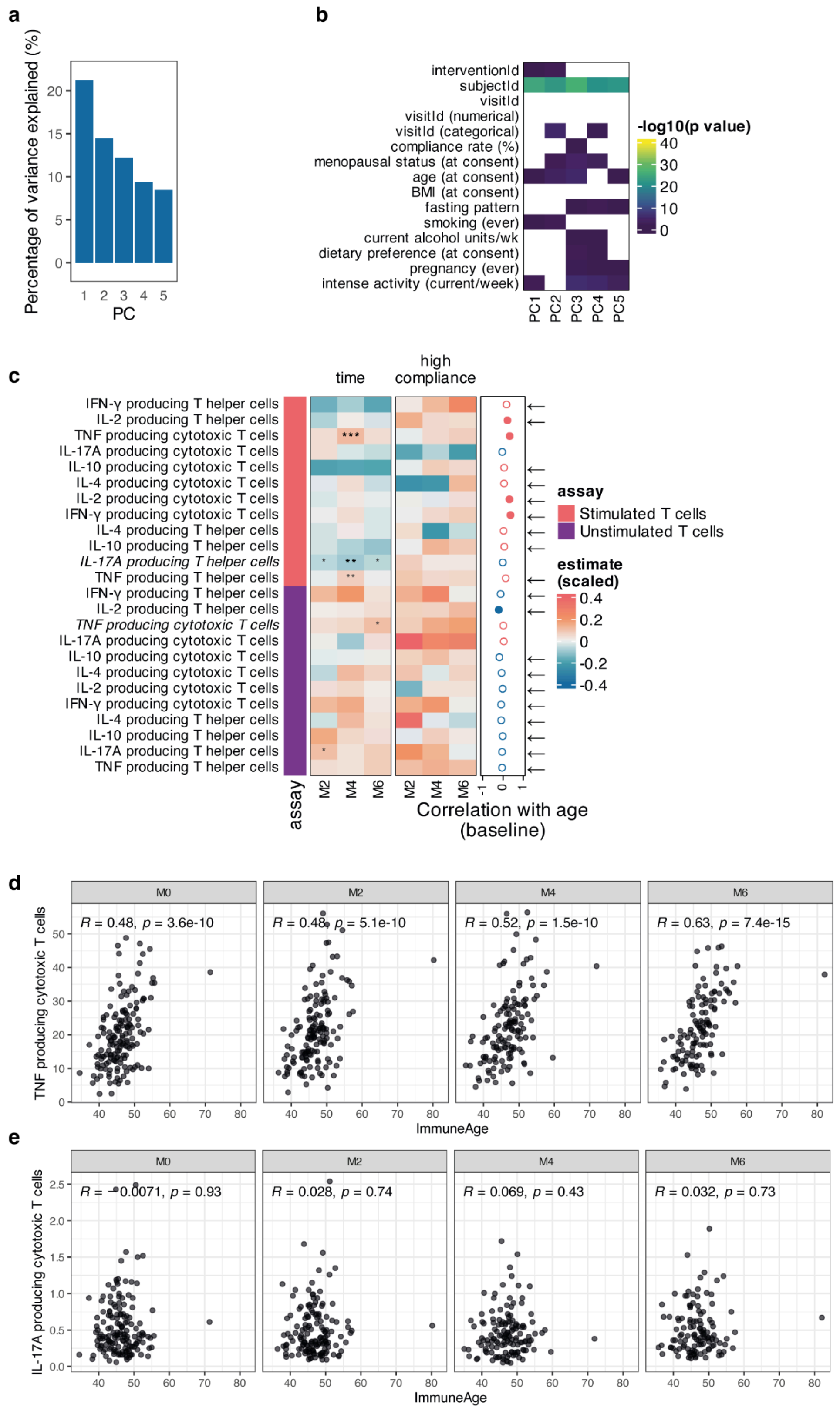


**Extended Data Figure 7. T cell cytokine production is altered over the course of intermittent fasting.**

**a** Scree plot showing the percentage of variance explained by the first five principal components in cytokine production profiling. **b** Heatmap of p values for associations of principal components with participant and study characteristics. **c** Estimates from linear-mixed effects models impact of time (value ~ age at consent + bmi at consent + visitId + (1|subjectId)) and interaction between time and high compliance (value ~ age at consent + bmi at consent + visitId*compliance + (1|subjectId)). **d**

Correlation of TNF-producting cytotoxic T cell fraction (after stimulation) with ImmuneAge at baseline and follow-up visits. **e** Correlation of IL-17A-producting cytotoxic T cell fraction (after stimulation) with ImmuneAge at baseline and follow-up visits.

For d and e, *, **, *** indicate $p < 0.05$, $< 0.01$, and $< 0.001$, respectively. Values that remain significant after FDR correction are indicated in bold; row labels of significant features after FDRare shown in bold. Values that remain significant after permutation testing correction are indicated in grey; row labels of significant features at month 6 after permutation testing are shown in italic. Correlation with age at baseline values is indicated on the right hand side (positive or negative in red and blue, respectively). Significant correlations at $p<0.05$ are indicated with filled circles. Populations that go into the opposite direction as ageing over time in the study are indicated with arrows.

Correlations in **d** and **e** indicate Spearman’s correlation coefficient.

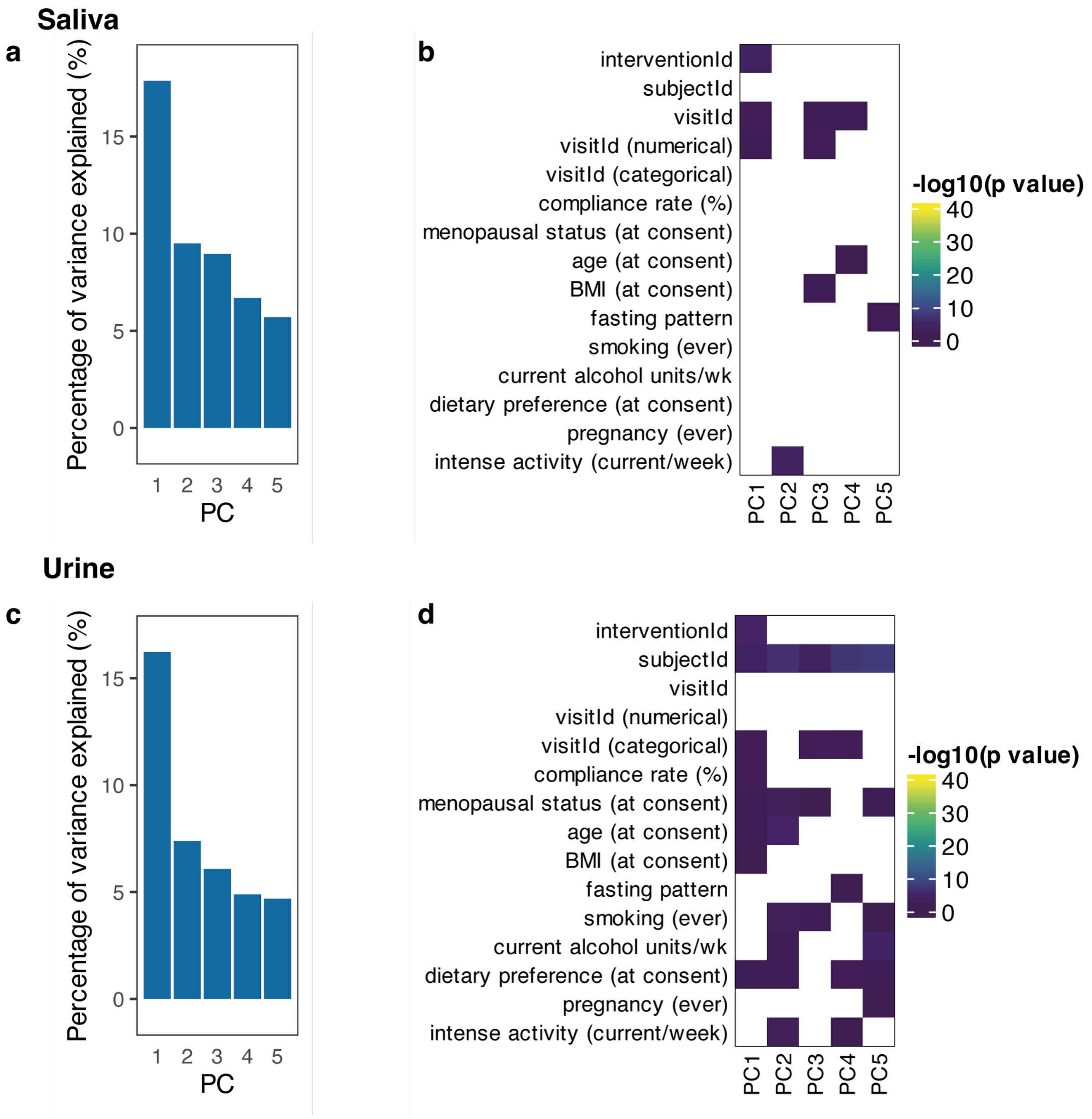


**Extended Data Figure 8. Principal component analysis on saliva and urine metabolome data.**

**a** Scree diagram of first five principal components in saliva metabolome PCA analysis. **b** Heatmap of analysis of features with the first five principal components in saliva metabolome data. **c** Scree diagram of first five principal components in urine metabolome PCA analysis. **d** Heatmap of analysis of features with the first five principal components in urine metabolome data.

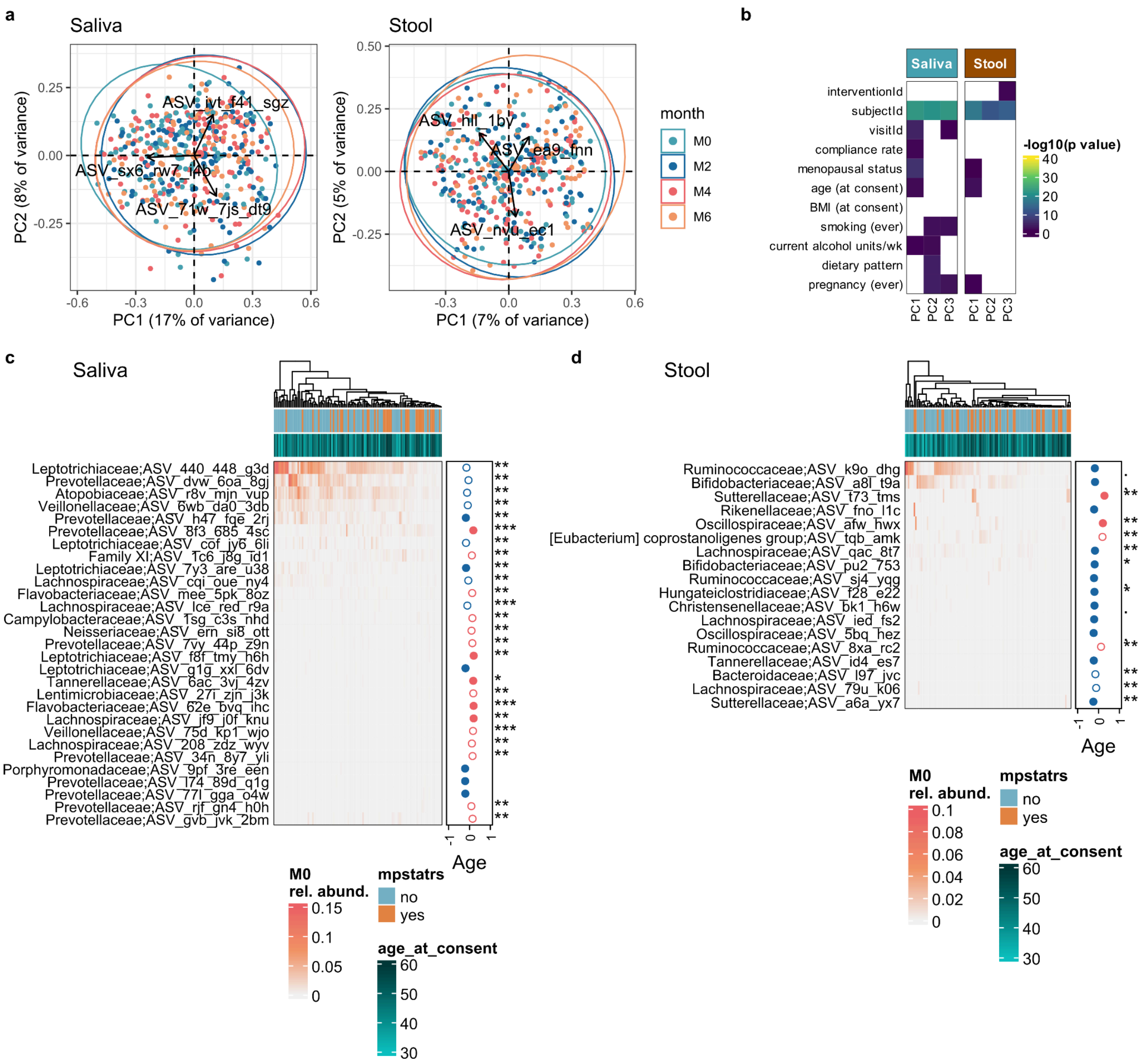


**Extended Data Figure 9. PCA on microbial ASVs and ASVs associated with age and menopausal status.**

**a** Biplots summarise the principal component analyses done on Hellinger transformed count data at the microbial ASV-level for the saliva and stool sample sets. Arrows depict the strength and direction of the top three features contributing to the overall variance. **b** Heatmap showing the significance of the first three principal components with recorded covariates. **c-d** Microbial ASVs significantly associated (Kendall correlations and Kruskal Wallis test, p<0.01 ) with age or menopausal status in the saliva and stool samples, respectively. Full circles indicate p value<0.01, for the association with age. *, **, *** denote p<0.05, <0.01, and <0.001, respectively, for the association with menopausal status.

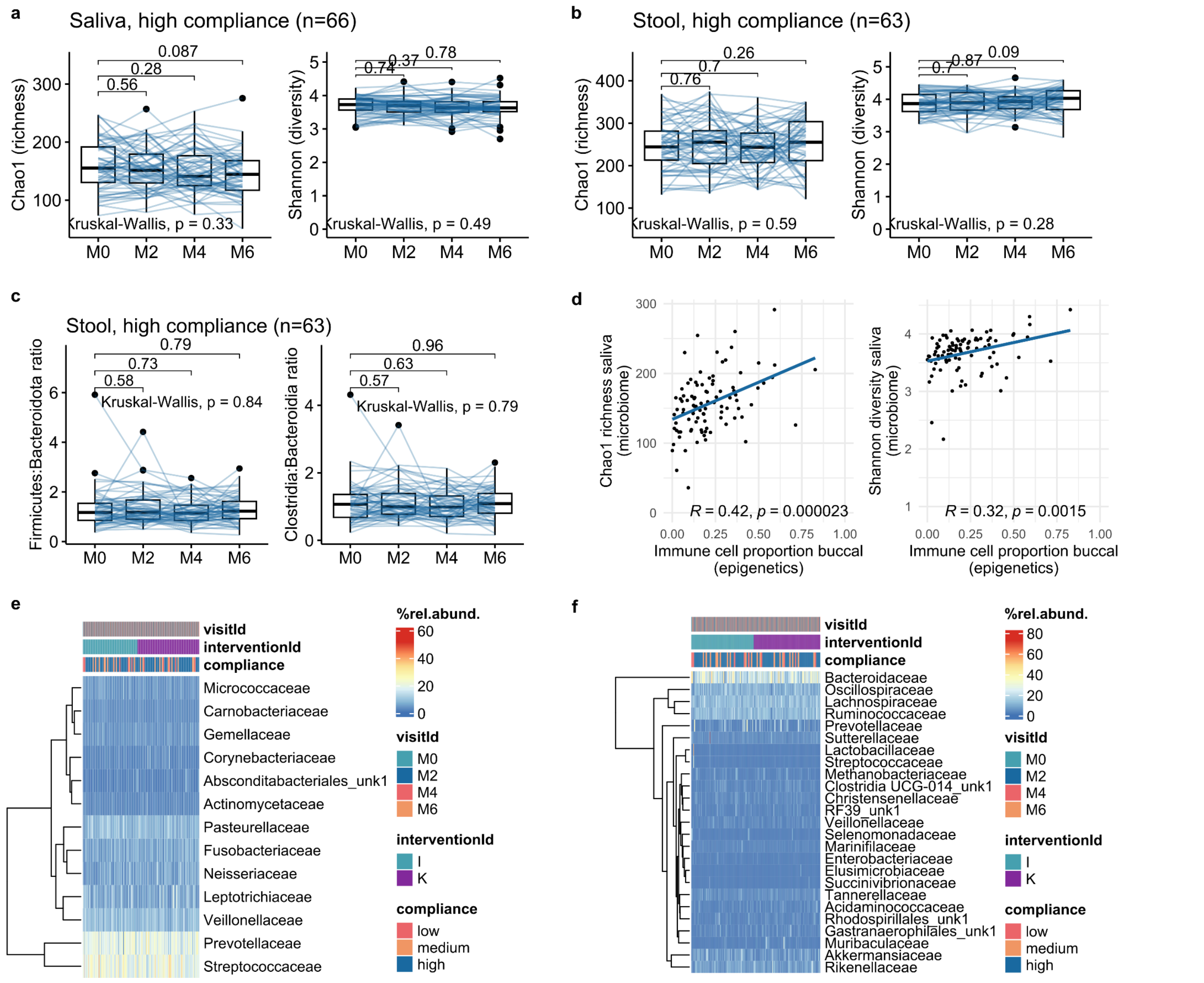


**Extended Data Figure 10. Alpha-diversity indices and family-level taxonomic composition of the microbiomes.**

**a-b** Chao1 and Shannon indices at different visits calculated for individuals highly compliant to the intermittent fasting intervention in saliva and stool, respectively. Boxplots indicate median and interquartile range while whiskers indicate values +/- 1.5 interquartile range. Individual data points are shown where possible. **c** Firmicutes:Bacteroidota and Clostridia:Bacteroidia ratios in gut microbiomes from individuals highly compliant to the intermittent fasting intervention. **d** Association of the Chao1 and Shannon indices in the saliva samples with estimated immune cell proportions in buccal epigenomes. Pearson correlation coefficient and associated p values are depicted.

**e-f** Microbial families that reached minimal 10% relative abundance in any of the saliva and stool samples, respectively.

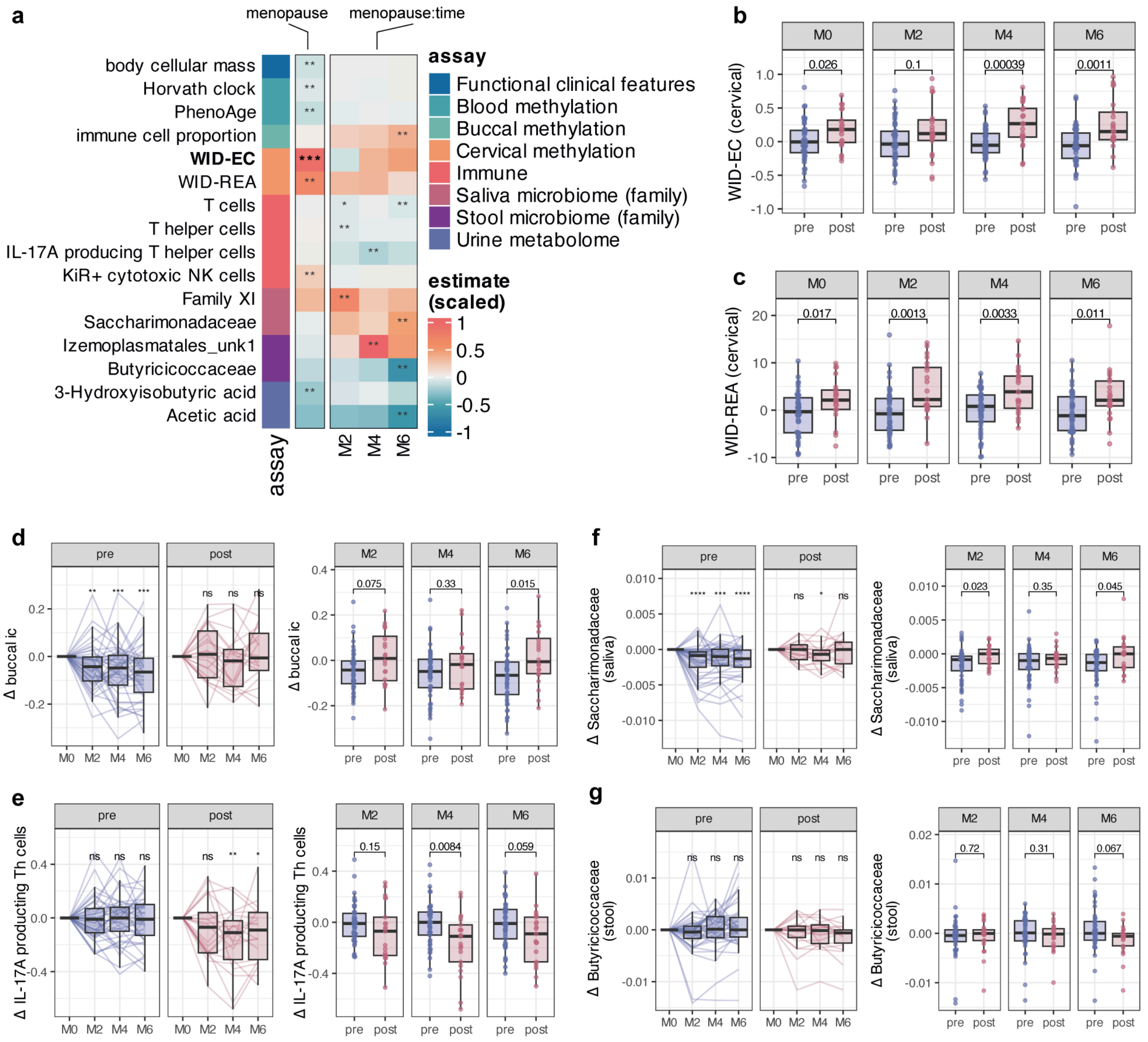


**Extended Data Figure 11. Features associated with menopause and treatment interaction with menopause.**

**a** Heatmap shows features across the difference omes that were significantly modulated by menopause or the interaction between time and menopause in two separate linear-mixed effects models (model 1: value ~ age at consent + bmi at consent + mpstatrs + visitId + (1|subjectId), model 2: value ~ age at consent + bmi at consent + mpstatrs:visitId + (1|subjectId)) in high compliance individuals only. FDR corrected p values are shown in bold. (**Extended Data Tables 9, 10**). *, **, *** indicate p<0.05, <0.01, and <0.001, respectively. **b-c** Significant differences between pre- and post-menopausal women for features picked up by model 1 were verified at each time point with paired two-sided Wilcoxon tests, with WID-EC and WID-REA shown here as two examples. **d-g** Significant interactions between menopausal status and intermittent fasting (time) picked up by model 2 were verified with paired two-sided Wilcoxon tests, comparing absolute values at M2-M6 to baseline in the pre- and post-menopausal groups, respectively, or comparing change from baseline in pre- and post-menopausal women at each time point using unpaired two-sided Wilcoxon tests. *, **, *** indicate p<0.05, <0.01, and <0.001, respectively.

Boxplots indicate median and interquartile range while whiskers indicate values +/- 1.5 interquartile range. Individual data points are shown where possible.

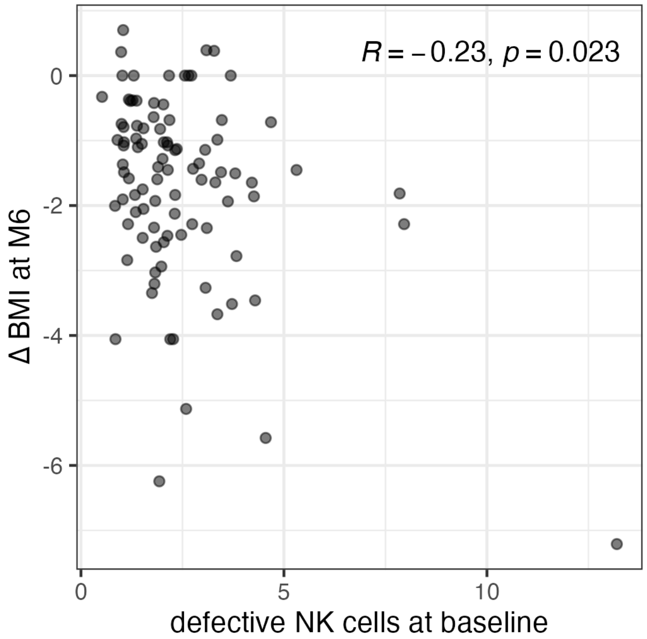


**Extended Data Figure 12. Defective NK cells at baseline predict ∆BMI at month 6.**
Correlation is Spearman's correlation coefficient.

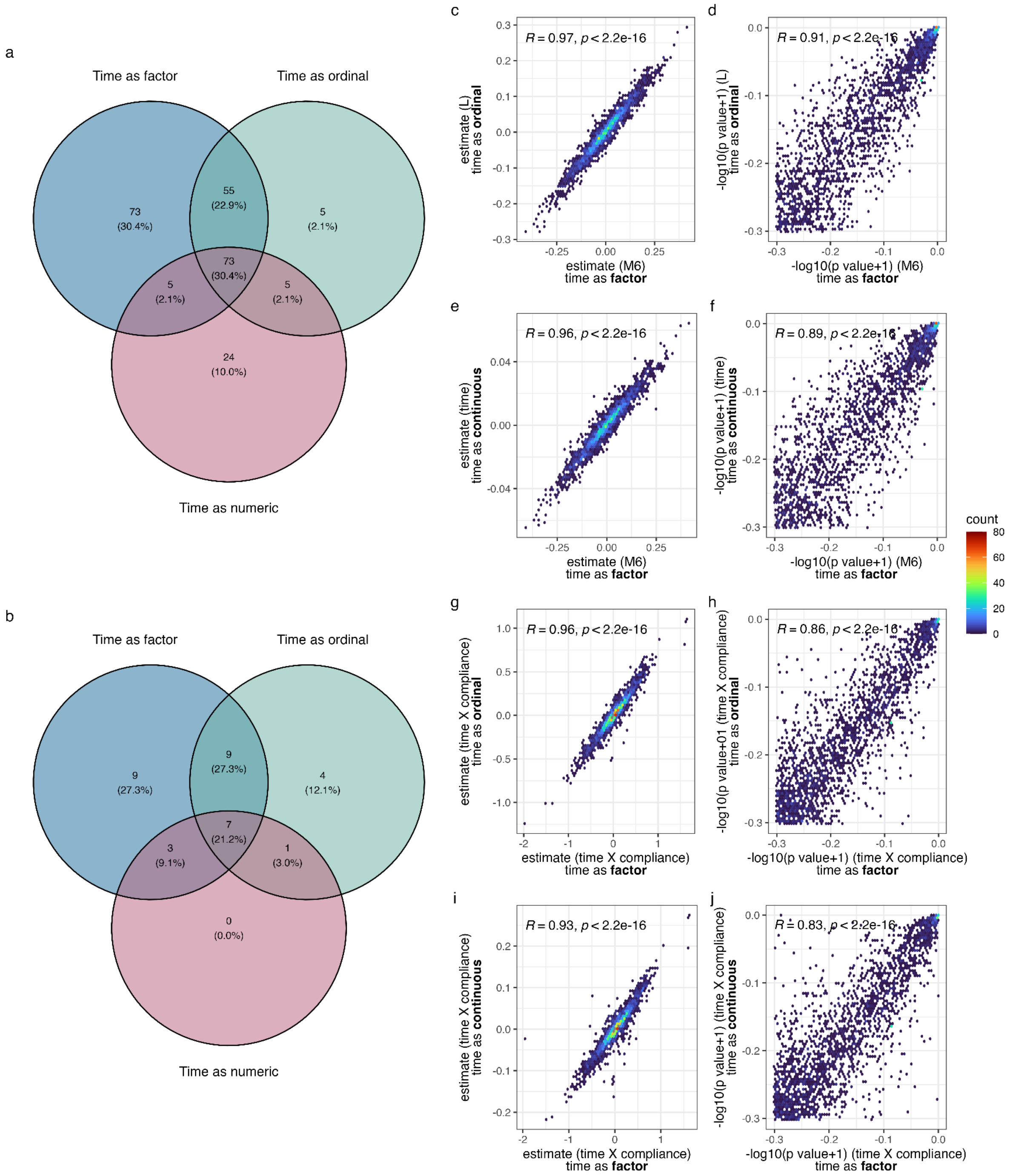


**Extended Data Figure 13. Sensitivity analysis of models with regards to coding the time variable as a categorical (factor), ordinal, or numeric for basic models (a, c-f) or interaction models (b, g-j).**

**a** Venn diagram depicting overlap of significant features at within-ome FDR < 0.05 in each basic model, testing the impact of time on the feature. **b** Venn diagram depicting overlap of significant features at within-ome FDR < 0.05 in each interaction model, testing the impact of the interaction of time and high compliance on the feature. **c** Correlation of effect estimates and **d** -log10(p values+1) for visitId month 6 (categorical/factor model) versus the linear estimate in the basic model. **e** Correlation of effect estimates and **f** -log10(p values+1) for visitId month 6 (categorical/factor model) versus the continuous time estimate in the basic model. **g** Correlation of effect estimates and **h** -log10(p values+1) for visitId month 6 (categorical/factor model) versus the ordinal estimate in the interaction model. **i** Correlation of effect estimates and **j** -log10(p values+1)

for visitId month 6 (categorical/factor model) versus the continuous estimate in the interaction model.
R denotes Pearson’s R (c, e, g, i) or Spearman’s rho (d, f, h, j), respectively.

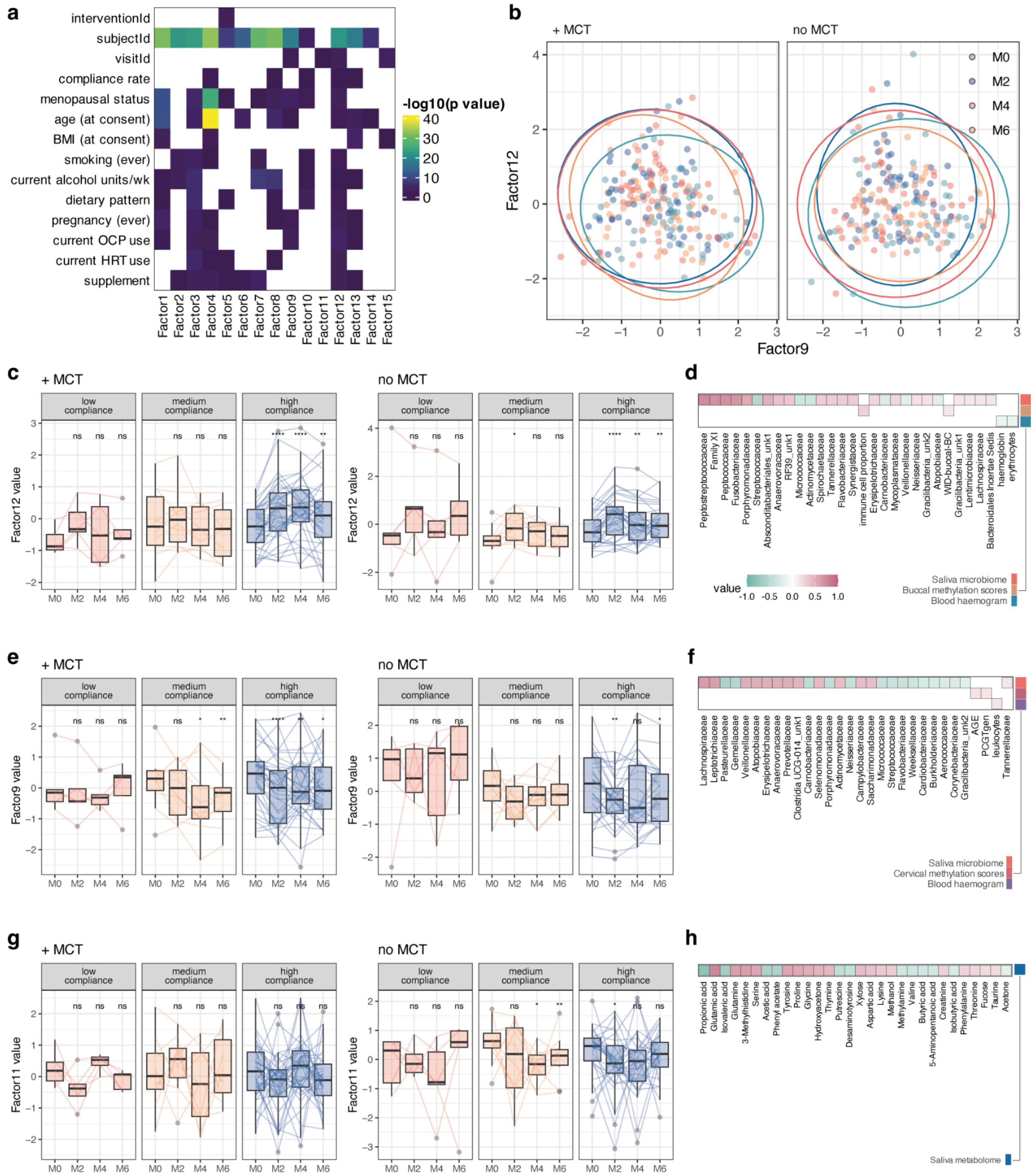

**Extended Data Figure 14. Integrative omic MEFISTO analyses.**

**a** Biplots depict the factor values for the top 2 latent factors found by MEFISTO in the two groups of participants (+/- MCT) using the epigenomes (composite methylation score and immune age), metabolomes, microbiomes, T cell and white blood cell stainings, and blood haemogram. The mean of the variances explained in each omic dataset by the first and second factors is given on the x, and y axes, and samples are coloured by visitId. **b** Heatmap showing the significance of the eight detected latent factors with recorded covariates. **c** Grouped bar plots give a breakdown of the different compliance and intervention groups for the latent factors significantly associated with visitId. Significance changes over time for each subgroup was verified with paired two-sided Wilcoxon tests (ns, *, **, ***, **** denote $p > 0.05$, $p \leq 0.05$, $p \leq 0.01$, $p \leq 0.001$, $p \leq 0.0001$, respectively). **d** Weights of the top 30 features associated with latent factor 1. Boxplots indicate median and interquartile range while whiskers indicate values +/- 1.5 interquartile range. Individual data points are shown where possible.

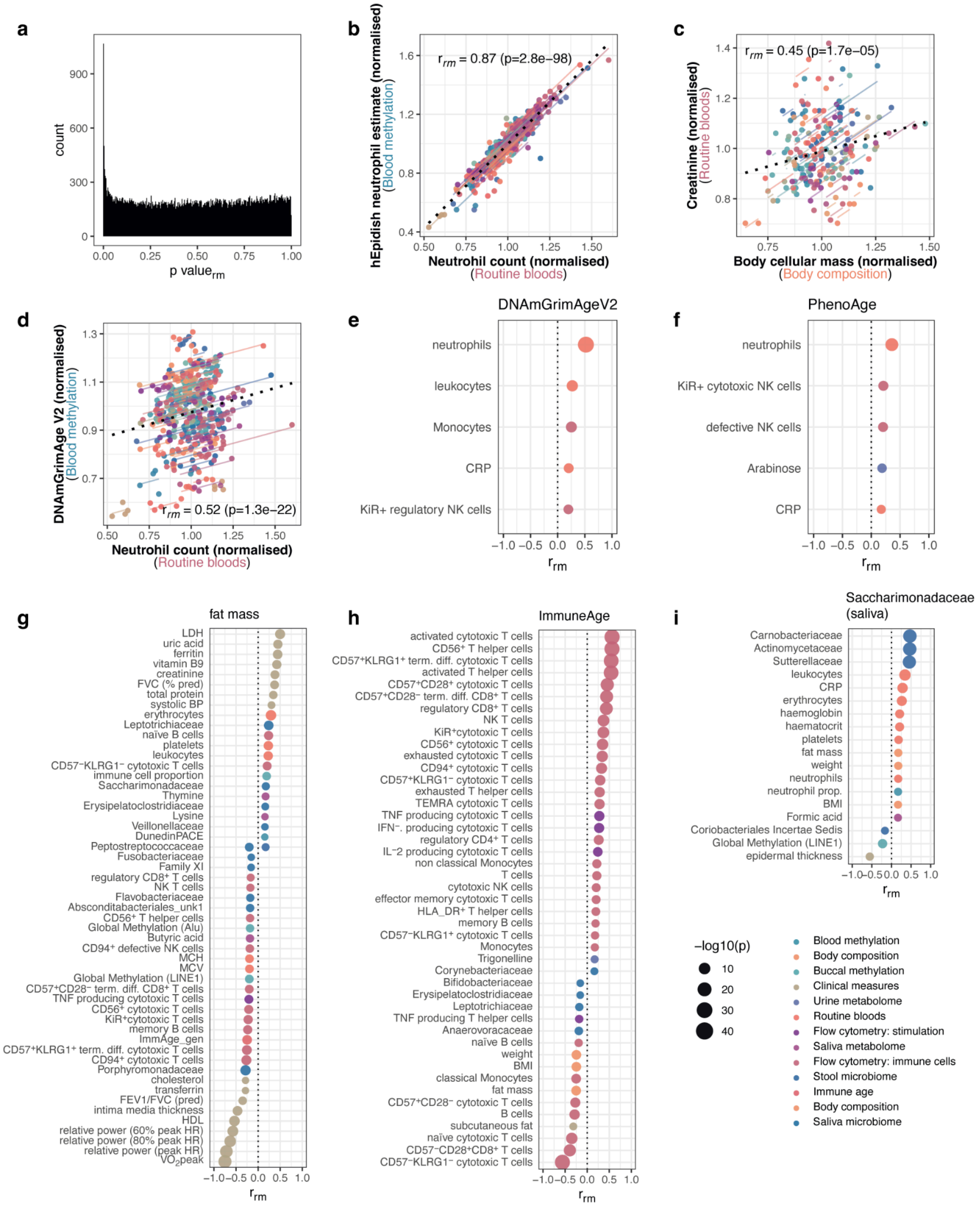


**Extended Data Figure 15. Repeated measures correlation results and examples for ageing--related biomarkers and most connected features.**

**a** p value histogram of the repeated measures (rm) correlation analysis. **b** Repeated measures correlation of estimated neutrophil proportion from blood methylation data and true neutrophil counts on routine blood tests. **c** Repeated measures correlation of creatinine values and body cellular mass. **d** Repeated measures correlation of AgeAccelGrim and neutrophil counts. **e** Repeated measures correlation of significantly correlated features with GrimAge (V2), **f** PhenoAge, **g** fat mass and **h** Immune Age, and **i** saliva Saccharimonadaceae. Colours indicate type of assay while size of the dots indicates the negative logarithm of the p value.

Colours in b-d indicate individual samples with model fit lines while dotted black line indicates overall correlation. The repeated measures correlation value $r_{rm}$ and p values (uncorrected) are shown; only p values that remained significant after FDR correction ($p < 0.05$) are retained.

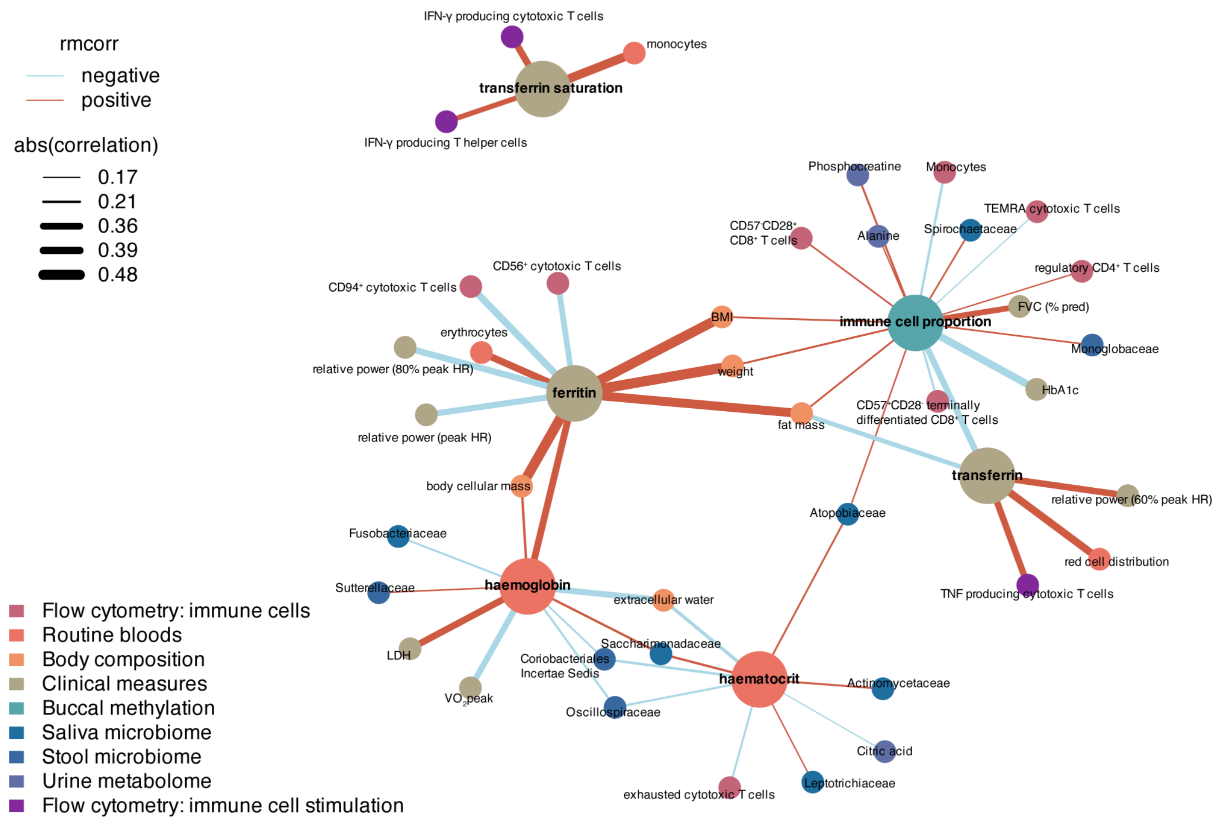


**Extended Data Figure 16. Repeated measures correlation network of iron-related metabolism.**

Correlations are extracted from repeated measures correlation (Extended Data Table 12), with only immediate first connections of transferrin, immune cell proportion, ferritin, haematocrit, haemoglobin, and transferrin and their first connections (|rmcorr|>0.6) shown in the network.

### Supplementary Data

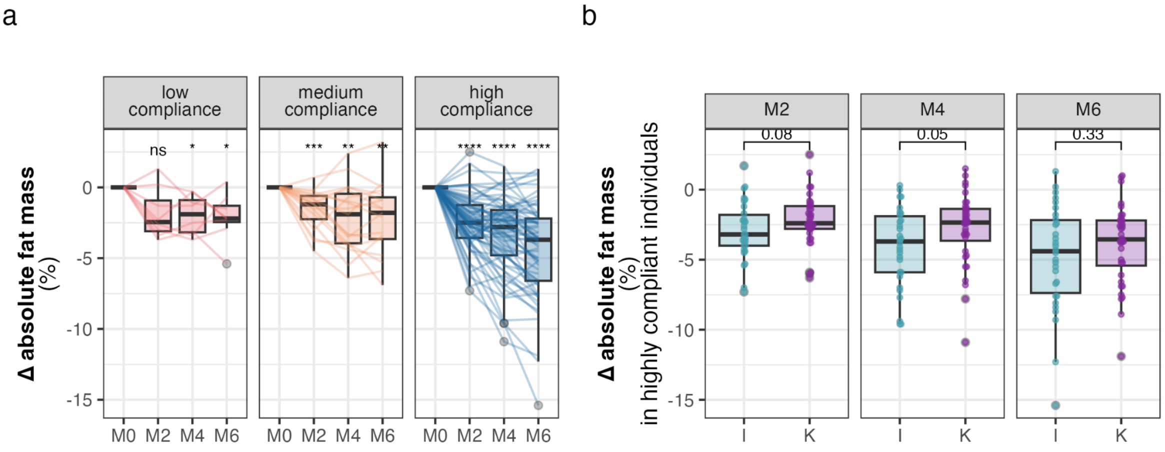


**Supplementary Figure 1. Data for fat mass illustrating impact of compliance and intervention, as summarised in Figure 2k.**
**a** Change in fat mass from baseline by compliance with intermittent fasting. p values derived from paired two-sided Wilcoxon tests compared to baseline values. **b** Change in fat mass by interventionId and time in highly compliant individuals only. p values derived from unpaired two-sided Wilcoxon test comparing I and K group.
**Abbreviations:** I, intermittent fasting only group. K, intermittent fasting plus ketogenic supplement group.
Boxplots indicate median and interquartile range while whiskers indicate values +/- 1.5 interquartile range. Individual data points are shown where possible.
